# Histone H3 E50K mutation confers oncogenic activity and supports an EMT phenotype

**DOI:** 10.1101/2023.10.11.561775

**Authors:** Kirti Sad, Celina Y Jones, Dorelle V Fawwal, Emily J Hill, Katie Skinner, Miranda Adams, Severin Lustenberger, Richard S Lee, Sandhya V Lohano, Satvik R Elayavalli, Jonathan Farhi, Laramie D Lemon, Milo B Fasken, Andrew L Hong, Steven A Sloan, Anita H Corbett, Jennifer M Spangle

## Abstract

Sequencing of human patient tumors has identified recurrent missense mutations in genes encoding core histones. We report that mutations that convert histone H3 amino acid 50 from a glutamate to a lysine (H3E50K) support an oncogenic phenotype in human cells. Expression of H3E50K is sufficient to transform human cells as evidenced by a dramatic increase in cell migration and invasion, and a statistically significant increase in proliferation and clonogenicity. H3E50K also increases the invasive phenotype in the context of co-occurring BRAF mutations, which are present in patient tumors characterized by H3E50K. H3E50 lies on the globular domain surface in a region that contacts H4 within the nucleosome. We find that H3E50K perturbs proximal H3 post-translational modifications globally and dysregulates gene expression, activating the epithelial to mesenchymal transition. Functional studies using *S. cerevisiae* reveal that, while yeast cells that express H3E50K as the sole copy of histone H3 show sensitivity to cellular stressors, including caffeine, H3E50K cells display some genetic interactions that are distinct from the characterized H3K36M oncohistone yeast model. Taken together, these data suggest that additional histone H3 mutations have the potential to be oncogenic drivers and function through distinct mechanisms that dysregulate gene expression.

**Summary:**

- Recurrent missense mutation that alter histone H3E50 were identified in patient tumors
- H3E50K expression dysregulates global H3 post-translational modification, gene expression and may govern a transcriptional program associated with metastatic phenotypes
- *hht2-E50K* yeast cells exhibit growth defects in the presence of DNA damaging agents

## Introduction

Routine patient tumor sequencing has unmasked recurrent somatic genomic alterations associated with cancer initiation, progression, and therapeutic response (1–4). Collectively, these studies have led to the identification and subsequent characterization of novel tumor suppressors and oncogenes, many of which are involved in transcriptional regulation and include epigenetic enzymes or complexes involved in chromatin reading, writing, and erasure (e.g., NSD1, DNMT3A, IDH1, and many others) (1,2,5,6). These and other functional studies provide evidence that appropriate chromatin packaging, modification, and organization are key contributors to the regulation of gene expression, and show that the role of chromatin dynamics extends into other essential processes that are frequently perturbed in cancer, such as DNA repair. Taken together, these studies show that various mechanisms that perturb chromatic structure can contribute to oncogenesis.

Chromatin is organized into repeating nucleosome subunits, wherein each nucleosome is comprised of 146-147bp of DNA, which wraps around a hetero-octamer of the histone proteins H2A, H2B, H3, and H4. Nucleosome-incorporated histones are subject to post-translational modifications (PTMs) directed by chromatin modifying enzymes, which enable context dependent epigenetic regulation to inform gene expression. Studies demonstrate that histone genes harbor genomic alterations in cancers and that oncogenic-driver variants of histone proteins dysregulate gene expression to contribute to an oncogenic phenotype (7,8), leading to the term “oncohistone.” Such oncohistone variants have been most extensively characterized in histone H3 (9–11). Although the human genome encodes 15 histone H3 genes, for most characterized oncohistones, mutation of a single H3 allele is sufficient for the development of oncohistone-driven cancers, demonstrating a dominant effect of the oncohistone mutant protein.

While numerous mutations in histone genes have been identified (8,12), only a few oncohistones have been functionally characterized. The H3K27M amino acid change occurs in at least 80% of pediatric diffuse intrinsic pontine gliomas (DIPGs) and has been reported with a lower frequency of 15-60% in adult diffuse midline gliomas (13,14). H3K36M has been identified in several cancer types, including chondroblastoma and squamous cell cancers of the head and neck (SCCHN)(15), while H3G34V/R and H3G34W/L predominantly occur in non-brainstem gliomas and bone cancers, respectively (8,16). More recent characterization of histone H2B mutations suggests that H2BE76K is found across cancer types, with a small preference for bladder and cervix (17). Functional studies support the hypothesis that oncohistones alter the chromatin landscape and perturb gene expression through distinct mechanisms. While H3K27 and H3K36 harbor PTMs that contribute to chromatin accessibility (7), mutation that converts either residue to a methionine impairs the ability of H3 to support PTMs, including methylation that can impact gene expression on a larger scale. H3K27M mutant histones reduce H3K27 methylation in *cis* and in *trans* by impairing polycomb repressive complex 2 (PRC2) recruitment to the genome, thereby reducing H3K27me3 genomic spread (18,19). The subsequent reduction in deposition of the transcriptionally repressive H3K27me3 PTM supports expression of development-associated genes, causing a de-differentiated and aggressive tumor type (18,20). In contrast, expression of H3K36M mutant histones dominantly impairs the recruitment and function of H3K36 methyltransferases including NSD1 and SETD2 (21,22). Because nucleosomes characterized by H3K36me3 are poor substrates for PRC2, loss of gene body H3K36me3 supports the spread of H3K27me3, which can aberrantly repress gene expression (10). H3G34 histones do not directly harbor PTMs, but alter proximal PTMs in *cis,* including H3K27 and H3K36 methylation (23,24). In contrast, the oncogenic H2BE76K mutation destabilizes the nucleosome core, leading to a more open chromatin conformation and enhanced gene expression (17). While other oncogenic histone mutations have been proposed, to date, most remain uncharacterized.

Recent preclinical studies have focused on leveraging the gene expression changes defined in oncohistone expressing human cancers to identify viable therapeutic targets. While H3K27M DIPGs do not induce extensive global gene expression changes, H3K27M tumors show dysregulation of a subset of genes with oncogenic activity, some of which may be therapeutically actionable (18). Gliomas, including H3K27M DIPG, are characterized by upregulation of the dopamine receptor gene *DRD2* (25) and increased PI3K/AKT and RAS/MAPK signaling (26). Initial proof-of-concept clinical trials using the DRD2 antagonist and mitochondrial ClpP agonist ONC201 in H3K27M-mutant diffuse midline glioma patients with recurrent disease demonstrate an improved patient quality of life and extended progression-free survival (25,27). Larger clinical trials have found that ONC201 treatment nearly doubles progression-free survival and overall survival in both nonrecurrent and recurrent diffuse midline gliomas (28). Other proteins, including EZH2 and STAT3, have been proposed as viable targets for the treatment of H3K27M-mutant cancers (29,30). These preclinical and clinical studies support the hypothesis that many oncohistone-driven cancers may be therapeutically actionable. Moreover, given that some oncohistones change gene expression and alter chromatin dynamics (e.g., H2BE76K), the opportunities for therapeutic treatments and therapy combinations may be extensive.

A challenge with understanding how the amino acid changes that occur in oncohistones alter histone function in the context of chromatin is that histone genes are present in multiple copies, creating a situation where a single oncohistone protein is present in the background of wildtype histones. One approach that has been utilized to understand how specific amino acid changes alter histone function in the context of chromatin has been to employ model systems (31–34). This approach benefits from the fact that the histone proteins are some of the most evolutionarily conserved proteins (35) and the fact that simple model systems such as yeast have far fewer copies of histone genes than higher eukaryotes. For example, while there are 15 histone H3 genes in humans, the budding yeast *Saccharomyces cerevisiae* has only two histone H3 genes, *HHT1* and *HHT2*. In a system such as budding yeast, one histone gene can be readily edited to model an oncohistone while either leaving the other histone gene intact, to assess whether the oncohistone protein confers dominant phenotypes similar to the situation in patients, or the second histone gene can be easily deleted to create cells that express the oncohistone as the sole histone present. In this latter scenario, studies can be performed to directly assess how the amino acid change present in the oncohistone alters the function of the histone protein. Studies in genetic model organisms can also take advantage of genetic strategies available in these systems to identify cellular pathways that are likely to be altered when oncohistones are present (34).

Here, we build upon publicly available data suggesting that H3E50K is a recurrent histone alteration in human cancer (7,8). Using a combined approach in untransformed human breast cells, *BRAF*-mutant human melanoma cancer cells, and a budding yeast model, we provide insight into this putative oncohistone H3E50K, which falls into a class of oncohistones that have the potential to generate novel sites for PTMs by introducing a surface-exposed lysine. We show that overexpression of H3E50K modestly increases cell proliferation and clonogenic growth in human breast cells, demonstrating that expression of H3E50K is sufficient to transform human cells. However, cells expressing H3E50K as the only putative oncogene or in the context of *BRAF-*mutant melanoma considerably enhance migratory and invasive potential, suggesting that H3E50K could be involved in cancer progression or other features common to metastatic disease. Cells expressing H3E50K exhibit changes to the H3 N-terminal tail and globular domain post-translational modification (PTM) profile, and dysregulation of gene expression. Compared to wildtype H3, gene signatures associated with the epithelial to mesenchymal transition (EMT) and oncogenic signaling pathways, such as KRAS and JAK/STAT, are increased in H3E50K breast cells. Genetic analyses exploiting a budding yeast model and comparing to an established H3K36M oncohistone yeast model (34) provide evidence to suggest that H3E50K may alter cell physiology through mechanisms that are distinct from H3K36M. These findings suggest that globular histone domain amino acid changes can drive oncogenic growth and could contribute to advanced stage cancer by initiation of the metastatic cascade.

## Materials and Methods

### In silico modeling

PDB files 5X7X (*H. sapiens*) (36) and 1ID3 (*S. cerevisiae*) (37) were used for all PyMOL imaging (Schrödinger). An overview of the human nucleosome structure was depicted in surface view, with DNA modeled in cartoon view. Higher-resolution images were obtained for the modeling of E50, K50, K50me3, K50ac, A50, and R50. For these images, the “mutagenesis” function was used to make amino acid substitutions. The lowest-numbered (highest-populated) state/rotamer for each amino acid residue was utilized to produce the images, given that it did not produce any clashes with surrounding amino acids. The PyMOL plugin “PyTMs” was utilized to model both trimethylation and acetylation of K50 (38). Interactions with nearby residues and DNA were measured in angstroms in the E50 and E50K states. The “ray” setting was used at 2400x2400 resolution to take all images, with “ray_shadow” set to off.

### Plasmids and cell lines

Wildtype *D. melanogaster* H3.3 (NCBI NM_001273153), which shares 100% amino acid identity with *H. sapiens H3.3,* was cloned into the pBabePuro IRES GFP plasmid (Addgene #14430). Wildtype *H. sapiens* H3.1 (NCBI NM_003531.2) was cloned into the pBabePuro plasmid (Addgene #1764). H3.3 and H3.1 mutations were introduced into the pBabePuro dH3.3-IRES GFP and pBabePuro plasmids, respectively, using site-directed mutagenesis (Agilent). Plasmids were verified by sequencing, packaged into retroviral particles, and used to generate stable HMEC cell lines. Authenticated HMEC cells were purchased from ATCC, engineered to express dominant negative p53 (39) and used for experiments within the first 20 passages. Cultures were checked for mycoplasma every three months (Lonza) and maintained at 37°C and 5% CO_2_ in DMEM/F12 media supplemented with 0.6% FBS, 0.01 µg/ml EGF, 10 µg/ml insulin, 0.025 µg/ml hydrocortisone, 1 ng/ml cholera toxin, 2.5 µg/ml Amphotericin B, and 1% Penicillin-Streptomycin.

### Human and yeast cell lysis and immunoblotting

Histones were acid extracted from human cells as described previously (40). In brief, cells were lysed on ice in Triton extraction buffer (TEB; PBS, 0.5% Triton X-100) supplemented with protease inhibitors [7.5 μM aprotinin, 0.5 mM leupeptin, 250 μM bestatin, 25 mM AEBSF-HCl]. Cell lysates were centrifuged at 6,500 x g at 4°C and histones were acid extracted from the resulting pellet with 1:1 TEB:0.8 M HCl. Histones were centrifuged at 4°C and the supernatant was precipitated with the addition of an equal volume of 50% TCA, then centrifuged at 12,000 x g at 4°C. Histones were washed one time in ice-cold 0.3 M HCl in acetone and two times ice-cold in 100% acetone before drying and resuspended in 20 mM Tris-HCl (pH 8.0), 0.4 N NaOH, supplemented with protease inhibitors [7.5 μM aprotinin, 0.5 mM leupeptin, 250 μM bestatin, 25 mM AEBSF-HCl]. Protein lysate concentration was determined by Bradford Assay (BioRad). 40 µg of acid-extracted proteins prepared in 1X LDA sample buffer (Invitrogen) containing 10% beta-mercaptoethanol were separated using SDS-PAGE. Proteins were transferred to nitrocellulose membranes and blocked in TBST + 5% milk. After primary and secondary antibody incubation, proteins of interest were visualized and quantified (Odyssey, Li-Cor).

Whole cell lysate was produced from human cells by lysing cells on ice in IPB (20 mM TrisHCl [pH 7.5], 150 mM NaCl, 5 mM MgCl_2_, 1% NP-40) supplemented with protease inhibitors [7.5 μM aprotinin, 0.5 mM leupeptin, 250 μM bestatin, 25 mM AEBSF-HCl]. Soluble proteins were separated via centrifugation at 15,900 x g for 10 min, transferred to a clean microcentrifuge tube, and protein concentration was determined by Bradford assay (BioRad). 30 µg of whole cell lysate prepared in 1X LDS sample buffer (Invitrogen) containing 10% beta-mercaptoethanol was separated using SDS-PAGE. Proteins were transferred to nitrocellulose membranes and blocked in TBST + 5% milk. After primary and secondary antibody incubation, proteins of interest were visualized and quantified (Odyssey, Li-Cor).

For yeast experiments, the indicated yeast strains were grown overnight at 30°C to saturation in 5 mL YEPD (yeast extract, peptone, dextrose) media. Cells were diluted in 100 mL YEPD to a starting OD_600_ = 0.1 and grown at 30°C to a final OD_600_ = 1.0. Cells were pelleted by centrifugation at 1,962 x *g* in 50 mL tubes, washed in ddH_2_O, transferred to 2 mL screwcap tubes, washed again in ddH_2_O, and pelleted by centrifugation at 900 x *g*. Pelleted cells were washed in Buffer 1 (1 M sorbitol, 50 mM Tris-HCl, pH 7.5, 5 mM MgCl_2_) and weighed. Samples were separated into two tubes to be <100 mg each. Cells were resuspended in 1 mL Buffer 1, and 5.4 μL 2-mercaptoethanol (14.3 M) was added to each. Samples were incubated on ice for 10 min. Cells were pelleted at 4°C at 900 x *g* and resuspended in 1 mL Buffer 1 plus 600 μg zymolyase 20T, then incubated at 35°C for 20 min with gentle agitation. Then, 0.7 mL Buffer 2 (1 M sorbitol, 50 mM MES, pH 6, 5 mM MgCl_2_) plus protease inhibitors (Thermo Scientific A32955) was added, and cells were pelleted at 1233 x *g* in 4°C. Samples were resuspended in 0.7 mL Buffer 3 (50 mM MES, pH 6, 75 mM KCl, 0.5 mM CaCl_2_, 0.1% NP-40) plus protease inhibitors, incubated on ice for 5 min, and pelleted at 12,175 x *g* at 4°C. Samples were resuspended in 0.7 mL Buffer 4 (10 mM MES pH=6, 430 mM NaCl) plus protease inhibitors and 0.5% IGEPAL CA-630, incubated on ice for 5 min, and pelleted at 14,489 x *g*. Finally, samples were resuspended in 0.7 mL Buffer 4 plus protease inhibitors, incubated on ice for 5 min, and pelleted at 17,005 x *g*. Pellets were resuspended in 120 μL 0.25 M HCl and spun in a rotor wheel at 4°C for at least 2 hr. Samples were pelleted at 14,489 x *g* in 4°C, and the supernatants were combined with 8 volumes of acetone and incubated overnight at -20C. Extracts were pelleted at room temperature at 1,962 x *g*, resuspended in acidified acetone (120 mM HCl in acetone), and pelleted 12,300 x *g*. Extracts were washed in acetone and pelleted. Extracted histones were air dried and resuspended in 50 μL ddH_2_O plus 1 μL NaOH. Protein lysate concentration was determined by Pierce BCA Protein Assay Kit (Life Technologies). Protein lysate samples (20-25 µg) in reducing sample buffer (50 mM Tris HCl, pH 6.8; 100 mM DTT; 2% SDS; 0.1% Bromophenol Blue; 10% Glycerol) were resolved on 4–20% Criterion™ TGX Stain-Free™ precast polyacrylamide gels (Bio-Rad). Protein lysate samples were transferred to nitrocellulose membranes (Bio-Rad) in Dunn Carbonate Buffer (10 mM NaHCO_3_, 3mM Na_2_CO_3_, pH 9.9, 20% methanol) at 22V for 90 min at room temperature, and the resulting membranes were blocked in TBST + 5% milk. After primary and secondary antibody incubation, proteins of interest were visualized and quantified using the ChemiDoc MP Imaging System (Bio-Rad).

For human and yeast studies, the primary antibodies used were as follows: H3 (Abcam 1791; 0.1 µg/ml), H3.3 (Abcam 17899; 1.71 µg/ml, TY1 (Diagenode C15200054; 2.2 µg/ml), HA (CST 6E2 1:1000), H3K4me3 (Abcam 8580; 1 µg/ml), H3K4me2 (Abcam 32356; 0.28 µg/ml), H3K4me1 (Abcam 8899; 2 µg/ml), H3K27me3 (CST 9733S; 1:1000), H3K27ac (CST 8173S; 1:1000), H3K36ac (Abcam 175038; 4 µg/ml), H3K36me3 (Abcam 9050; 1 µg/ml), H3K56ac (Invitrogen PA5-40101; 1.6 µg/ml), H3K79me3 (CST 4260S 1:1000), actin (Millipore MAB1501, 1:1000), N-Cadherin (CST 13116T, 142 ng/ml), E-Cadherin (CST 3195T, 54 ng/ml), and p53 (Santa Cruz DO-1; 100 ng/ml). Secondary antibodies include the fluorophore-conjugated goat anti-rabbit 680 IgG (LiCor, 925-68071; 1:10,000) and goat anti-mouse 800 IgG (LiCor, 926-32210; 1:1000).

### Quantitation of histone immunoblotting

The protein band intensities from immunoblots were quantitated using Image Lab software (Bio-Rad) and mean fold changes in protein levels were calculated in Microsoft Excel (Microsoft Corporation). The protein band intensity was normalized for each endogenous histone PTM level to the total endogenous H3 band. The mean fold change was further calculated related to the wildtype H3 sample for each PTM. The data were represented as an average fold change compared to wildtype with standard deviation (n=3). The mean fold changes in histone H3K36me3 levels in oncohistone mutant cells relative to the wildtype control were calculated from two immunoblots. H3K36me3 band intensity was first normalized to total histone H3 band intensity and then normalized to H3K36me3 intensity in wildtype cells. The mean fold changes in H3K36me3 levels in oncohistone mutant cells relative to the wildtype control were graphed in GraphPad Prism 8 (GraphPad Software, LLC) with standard error of the mean error bars.

### Transformation assays

To measure cell proliferation, 3,000 cells/well were seeded in a 96-well plate. Cells were fixed at 24, 48, 72, and 96 h post-seeding in 50% EtOH and 10% acetic acid, and crystal violet stained in 0.2% w/v crystal violet in 10% EtOH. Cells were destained in destaining solution (40% EtOH and 10% acetic acid) and crystal violet quantitated at OD595 (Biotek). To measure cell proliferation over an extended time period (15-18 d), 25,000 cells were seeded in a 24-well plate and cultured for 3 d with changing media every second day. After 3 d, the cells were trypsinized, counted, and 25,000 cells were re-seeded. At the end of the experiment, the total number of cells were calculated. To measure cell clonogenicity, 500 cells/well were seeded in a 6-well plate and incubated for a total of 15 d with media change every 4 d, after which cells were fixed and crystal violet stained as described above. Plates were imaged, destained, and crystal violet quantitated at OD_595_ (Biotek). To perform the cell confluence assay for population doubling, 2,000 cells were seeded in a 96-well plate (Corning 353072) in triplicate. The plate was scanned in Incucyte (Sartorius) at 37°C with 5% CO2 for 96 h using a time lapse system that imaged the cell confluency every 2 h. At the end of the experiment, the data was analyzed using Sartorius Rev2 software, and the growth curve was plotted based on the cell confluency over time.

### Wound closure assays

4 × 10^5^ cells were seeded in duplicate in 12-well plates. Once confluent, cells were scratched with a sterile p200 pipette tip. Images of the wound were collected at 0h, 5h, and 12h post scratch (Leica delimited). The rate of cell migration and percent wound closure was calculated using Image J.

### Transwell migration and invasion assays

For transwell migration assays, 1 × 10^5^ cells were seeded in duplicate in an 8.0 micron transwell 24-well plate insert (Corning 353097) in 100 µl of DMEM/F12 media without growth factors and serum and 600 µl full growth media in the bottom of the well. Non-migrated cells were removed from the upper transwell membrane with a cotton swab after 8 h. Transwell membranes were fixed in 70% EtOH for 10 min, allowed to dry for 10 min, followed by crystal violet staining in 0.2% w/v crystal violet in 10% EtOH. Excess crystal violet was removed from the upper transwell membrane with a cotton swab, and the inserts were washed in deionized water. After drying, the membrane was cut from the insert and mounted on glass slides. Images were collected (Leica delimited) and migrated cells were quantitated with Image J. For transwell invasion assays, 8.0-micron transwell 24-well plate inserts were coated with 100 µl of 1 mg/ml matrigel (Corning 254234). The plates coated with matrigel were incubated for 1 h at 37°C to solidify. 1 × 10^5^ cells were seeded in duplicate on the top of matrigel coated insert in 100 µl of DMEM/F12 media without growth factors and serum and 600 µl full growth media in the bottom of the well. Cells were incubated for 48 h for invasion and processed as described for the transwell migration assays.

### RNA isolation, sequencing, and analysis

RNA was isolated from HMECDD cells using the RNeasy Isolation Kit (Qiagen) according to the manufacturer’s instructions. Sequencing and analysis were performed as previously described (41). In brief, preparation of the RNA library and transcriptome sequencing was conducted by Novogene Co., LTD (Beijing, China). The resulting FASTQ files were aligned to the human reference genome (build GRChg38) and transcript abundance was quantified using salmon (Illumina DRAGEN). The DESeq2 package was used to identify differentially expressed transcripts between the H3.3E50K and wildtype H3.3 expressing HMECDD cells. Transcripts with adjusted *p*-value ≤ 0.05 and |log2(FoldChange)| ≥ ±1.5 were considered differentially expressed. Fast gene set enrichment analysis (FGSEA) of the pre-ranked gene list was performed using clusterProfiler and fgsea, using the msigdb hallmark gene set.

### S. cerevisiae strains and plasmids

Chemicals used for experiments with *Saccharomyces cerevisiae* were obtained from Sigma-Aldrich (St. Louis, MO), United States Biological (Swampscott, MA), or Fisher Scientific (Pittsburgh, PA) unless otherwise noted. All media were prepared by standard procedures (42). All DNA manipulations were performed according to standard procedures (43). *S. cerevisiae* strains and plasmids used in this study are listed in Table S2. The PCR- and homologous recombination-based system for generating targeted mutations in histone genes in budding yeast cells has been described (44). Strains to model oncohistones—*hht2-K36M hht1*ti (ACY2822), *hht2-E50K hht1*ti (ACY2972), *hht2-E50R hht1*ti (ACY2968), and *hht2-E50A hht1*ti (ACY2964), which harbor mutations at the codons encoding the 36th or 50th histone H3 residue at the endogenous *HHT2* gene—were generated using the parental *hht2Δ::URA3:TRP1* strain (yAAD165) and the strategy detailed previously (44,45). The endogenous *HHT1* gene in these oncohistone model strains was subsequently deleted and replaced via homologous recombination with a *kan*MX marker cassette. The isolation of the YEp352 genomic DNA plasmids from the suppressor screen – *HHT2_HHF2* (pAC4145), *SGV1* (pAC4132), *ESA1* (pAC4149), *TOS4_YLR184W* (pAC4150), *PHO92_WIP1_BCS1* (pAC4160) – and the YEp352 plasmids containing cloned *S. cerevisiae* genes – *HHT2* (pAC4201), *ESA1* (pAC4190), *TOS4* (pAC4196), *PHO92* (pAC4193), *SGV1* (pAC4208) – is described in (34).

### S. cerevisiae growth assays

To examine the growth of oncohistone model strains, wildtype (yADP127), *hht2-K36M hht1Δ* (ACY2822), *hht2-E50K hht1*ti (ACY2972), *hht2-E50R hht1*ti (ACY2968), and *hht2-E50A hht1*ti (ACY2964) strains were grown overnight at 30°C to saturation in 2 mL YEPD (yeast extract, peptone, dextrose) media. Cells were normalized to OD_600_ = 5, serially diluted in 10-fold dilutions, spotted on control YEPD media plates or YEPD media plates containing 18 μg/mL camptothecin, 6 μg/mL bleomycin, or 15 mM caffeine and grown at 30°C for 2-5 d. Cells were also grown on Ura-plates containing 2% glucose with or without 15 mM caffeine to ensure plasmid expression. To test the effect of the H3K36 mutant suppressor plasmids on the growth of H3E50 mutant strains in the presence of caffeine, wildtype (yADP127), *hht2-E50K hht1*ti (ACY2972), *hht2-E50R hht1*ti (ACY2968), and *hht2-K36M hht1Δ* (ACY2822) cells transformed with YEp352, *HHT2* (pAC4145), *ESA1* (pAC4149), *TOS4* (pAC4150), *PHO92* (pAC4160), or *SGV1* (pAC4132) plasmid were grown overnight at 30°C to saturation in 2 mL Ura-media containing 2% glucose. Plasmids in Figure 6 were isolated directly from the screen in (Lemon et al., G3 2022), and marked plasmids from Figure S5 were created by cloning. Cells were normalized to OD_600_ = 5 and serially diluted as previously described, spotted onto control YEPD media plates or YEPD media plates containing 15 mM caffeine, and grown at 30°C for 2-5 days. Cells were also spotted onto control Ura-plates or Ura-plates containing 15 mM caffeine.

### Statistical analysis

All experiments were performed in three independent experiments unless otherwise noted. Mean ± SD are reported unless otherwise noted. Statistical significance (p ≤ 0.05) of differences between two groups was determined by Student’s t test.

## Results

### Mutations that alter H3E50 are recurrent in human cancers

To date, most established histone H3 oncohistones function through direct perturbation of a known site of histone PTM (e.g., H3K27M and H3K36M/R), or an amino acid proximal to H3K36 (e.g., H3G34V/R/W/L) (7,8). These oncogenic changes can disrupt histone H3 PTMs in *cis* and/or in *trans*, imparting functional changes that result in transcriptional dysregulation. To determine whether additional histone mutations are associated with human cancers, we surveyed the COSMIC and cBioPortal publicly available adult human cancer datasets and identified recurrent mutations that alter H3E50 (46,47) (Figure 1). The E50 residue is located within the H3 globular domain (Figure 1A-C). H3E50 is surface accessible within the nucleosome and located in close proximity to DNA (Figure 1B, C). Mutations that alter H3E50 were identified in 37 adult cancer cases (Figure 1D). In contrast to H3K36M and H3K27M, which predominantly occur in one of the two *H3.3* (*H3-3A* or *H3-3B*) genes, H3E50 mutations primarily occur in *H3.1* genes, with cases also identified in *H3.2* and *H3.3* genes (Figure 1D, Figure S1A). With respect to the specific amino acid change introduced, H3E50K and H3E50D are the most common, although other changes were detected, including H3E50Q and H3E50*, with the latter introducing a premature stop codon (Figure 1E). Genomic alterations affecting H3E50 were detected most frequently in lung and breast cancers, although H3E50 mutations were detected in a variety of other cancer types, including melanoma (Figure 1F, Figure S1B). To extend the analysis of missense mutations that alter H3E50, we investigated whether genomic alterations that alter H3E50 co-occur with established oncogenic driver and/or tumor suppressor alterations in cancer. Using the cBioPortal and COSMIC databases, we identified co-occurring genomic alterations in tumor suppressors (e.g., *TP53, NF1* and *PTEN*) and oncogenes (e.g., *PIK3CA, BRAF, KRAS*). In fact, characterized mutations in *BRAF* (V600E), *PIK3CA* (E545K, H1047R), and *KRAS* (K117N) were found in patient tumors with H3E50 mutations (Figure 1G, Table S1). These results suggest that considering the functional consequences of oncohistone mutations in the context of these co-occurring mutations could be critical. To address whether H3E50 alterations are likely oncogenic driver events, we examine the allele frequency. H3E50 variant allele frequency ranged from 20-50% (Figure 1H), with an average H3E50K variant allele frequency of 27% (0.27± 0.15) in diploid tumor samples. The average H3E50 variant allele frequency is comparable to the known oncogenic driver mutation H2BE76K (17), and suggests that H3E50 alterations including H3E50K are likely acquired subclonal mutations and not germline mutations or polymorphisms.

**Figure 1.**
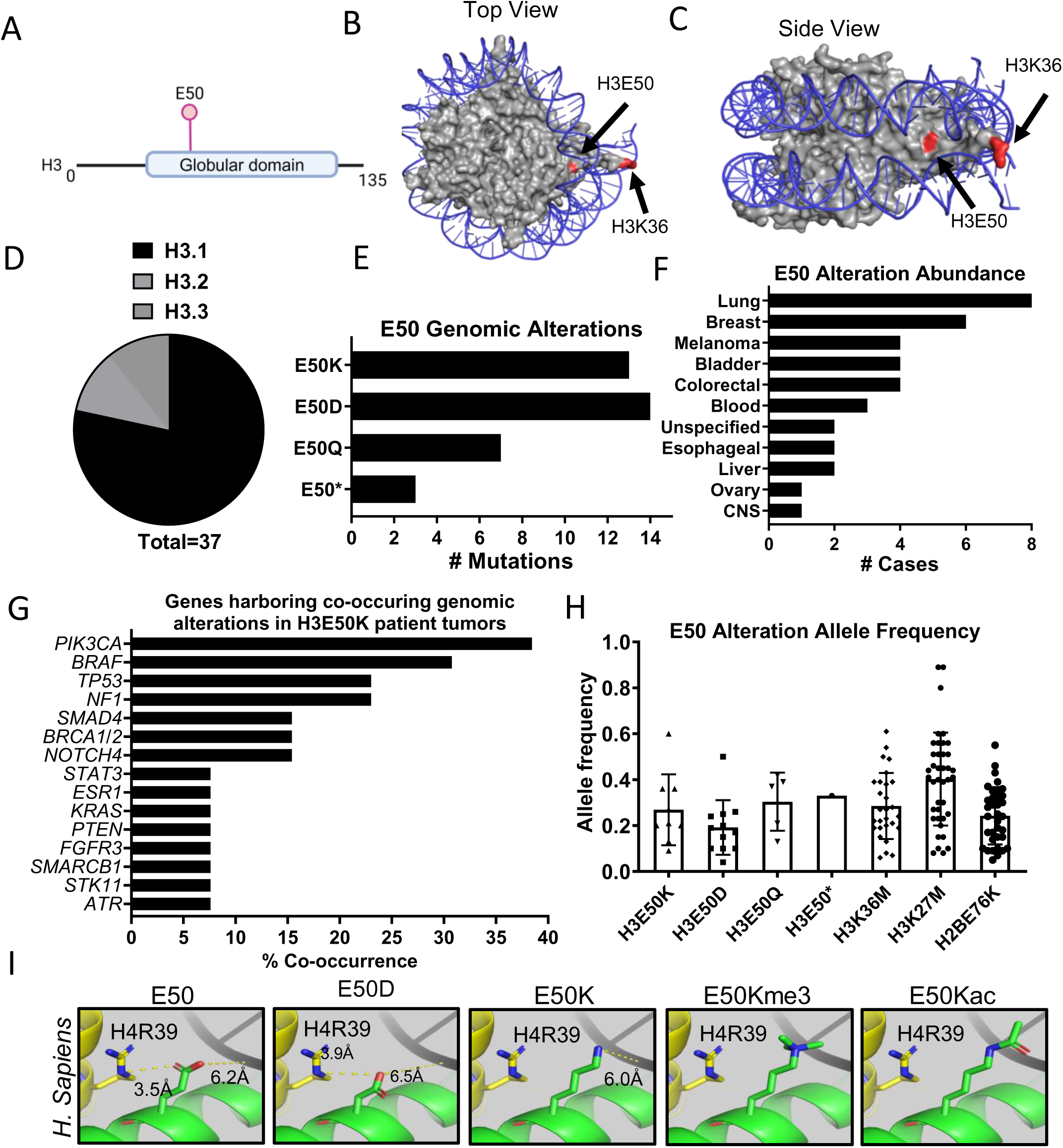
H3E50 mutation occurs in human cancers. (A) Schematic of H3.3 highlighting the E50 residue within the globular domain. (B) top and (C) side view of H3E50 and H3K36 within the nucleosome. Blue = DNA; Red = H3K36 and H3E50; Grey = remainder of amino acids encoding histone proteins H2A, H2B, H3, and H4, using PDB 5X7X (36). (D) Survey of E50 mutation abundance in histone H3 genes encoded in the human genome. (E) Identity of defined H3E50 mutations in human cancers. (F) Survey of H3E50 mutation abundance across human cancer types. (G) Percent co-occurring confirmed or probable oncogenic or tumor suppressor genomic alterations observed in patient tumors characterized by H3E50K mutation. (H) Boxplots showing variant allele frequency (VAF) of the H3E50 genomic alterations compared to known oncohistones H2BE76K, H3K27M, and H3K36M observed in patient tumors defined in cBioPortal. (I) *In-silico* modeling of human H3E50, H3D50, H3K50, and H3K50 harboring possible post-translational modifications, using PDB 5X7X (36).

As mutations that alter lysine residues, which can harbor PTMs, are common in previously studied oncohistones, we focused our efforts on examining the potential oncogenicity of H3E50K. *In silico* modeling demonstrates that H3E50 interacts with H4R39 (Figure 1I). Altering this residue to H3E50K creates a repulsion between the H3K50 R group and the H4R39 R group, which may locally open the nucleosome structure. In comparison, altering this residue to H3E50D does not change the interaction with H4R39. Should H3K50 support methylation or acetylation, a similar repulsive interaction with H4R39 could occur. Thus, a change in H3E50 could alter overall nucleosome structure or impact interaction with packaged DNA.

### H3E50K and H3E50D transform human cells and alter cellular growth

To examine whether H3E50 mutation exhibits properties consistent with oncogenic activity and is sufficient to transform human cells, we utilized the hTERT immortalized but untransformed human mammary epithelial cells (HMECs) that were previously engineered to express a dominant negative p53 mutant protein (39). Dominant-negative p53 expression in HMEC cells, or “HMECDD”, does not fully transform these cells; rather, these cells require expression of an oncogene in order to produce a transformed phenotype *in vitro* (39,48,49). We transduced HMECDD cells with vectors expressing a C-terminal TY1 epitope-tagged wildtype H3.3, H3.3E50K, or the established oncohistones, H3.3K27M or H3.3K36M (7), to produce stable, pooled HMECDD H3.3-TY1, H3.3E50K-TY1, H3.3K27M-TY1, and H3.3K36M-TY1 cell lines. Because H3.1 harbors genomic alterations H3.1E50K and H3.1E50D in human cancers (Figure 1D), we also transduced HMECDD cells with vectors expressing a C-terminal HA epitope-tagged wildtype H3.1, H3.1E50K, or H3.1E50D to produce stable, pooled HMECDD H3.1-HA, H3.1E50K-HA, and H3.1E50D-HA cell lines. To confirm expression of each histone variant, we acid extracted histones from these cells and analyzed ectopic H3.3 and H3.1 expression via the TY1 and HA tags, respectively. As shown in Figures 2A and 2B (Figure S2A, S2B), each of these ectopic histones are expressed at similar levels across this pool of cells, and ectopic histone expression does not change p53 expression (Figure S2C).

**Figure 2.**
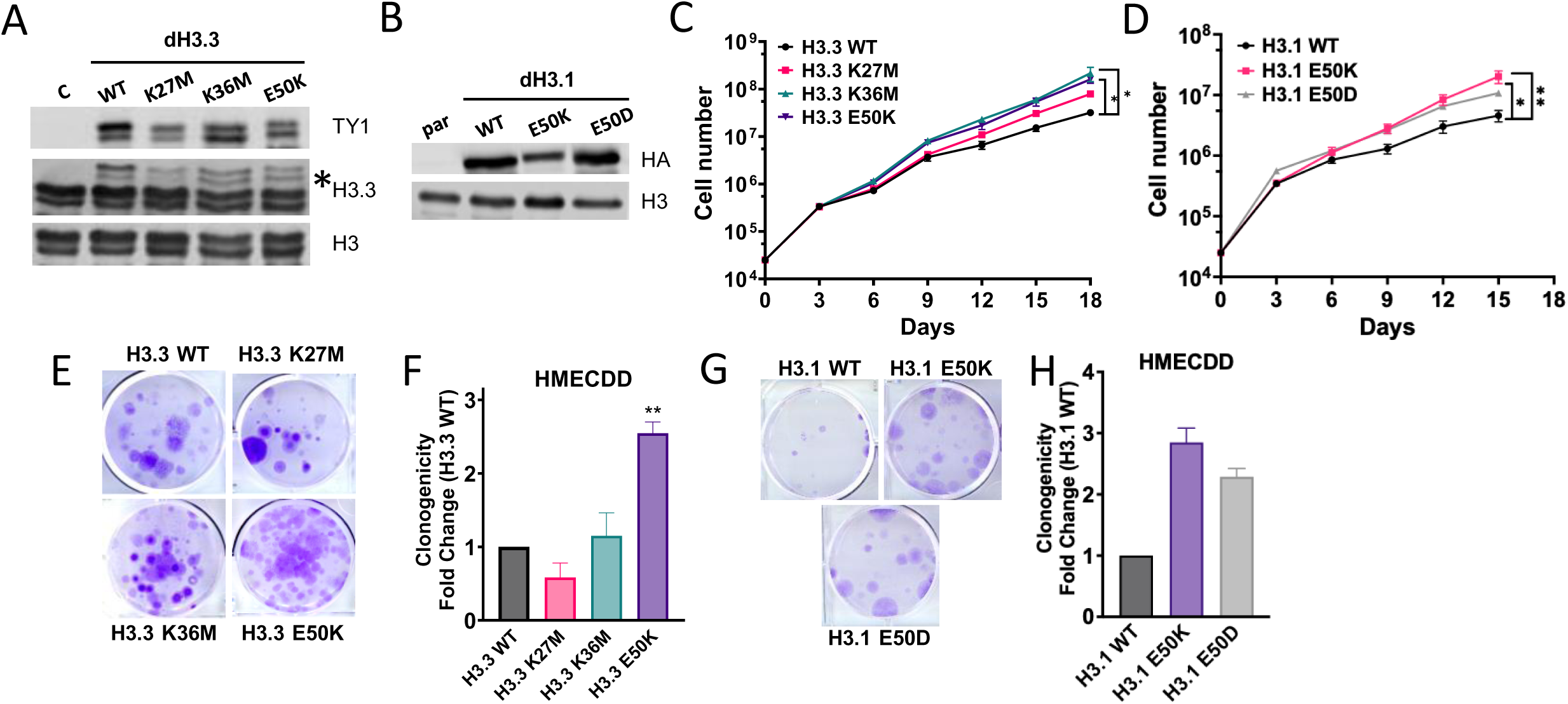
H3E50 variant expression increases cell proliferation and clonogenicity. HMECDD cells stably transduced with (A) pBabePuro H3.3-TY1, H3.3K27M-TY1, H3.3K36M, or H3.3E50K or (B) pBabePuro H3.1-HA, H3.1E50K-HA, or H3.1E50D and lysates acid extracted. Lysates were immunoblotted with the indicated antibodies. Representative images shown, *n = 3.* (C) and (D) Stable HMECDD cells expressing the indicated H3.3 mutant proteins were seeded and cell proliferation measured over the indicated time course, *n = 3.* (E) HMECDD cells stably transduced with the indicated H3.3 plasmids were seeded and cell clonogenicity measured after 15 d, representative images shown (F) Quantification of (E), *n = 3.* (G) Stable HMECDD cells expressing the indicated H3.1 mutant proteins were seeded and cell clonogenicity measured after 20 d, representative images shown. (H) Quantification of (G), *n = 4*.

To explore the functional consequences of H3E50K, we seeded this series of HMECDD cells and monitored cell proliferation over time. We found that H3.3E50K, H3.1E50K, and H3.1E50D expression significantly but modestly enhances HMECDD proliferation compared to wildtype histone H3 proteins (Figure S2D, S2E). We monitored cell growth over time using the Incucyte platform that measures cell confluence in real-time and found that H3.3E50K expression increases cell confluence over a 96 hour time course relative to wildtype H3.3 proteins, whereas H3.1E50K and H3.1E50D did not, suggesting a faster doubling time for H3.3E50K (Figures S2F, S2G). Upon examining cell proliferation for an extended time period of 15-18 days, we did find a significant increase in the cell proliferate rate for HMECDD cells expressing either H3.3E50K, H3.1E50K, or H3.1E50D, relative to wildtype H3.3 or H3.1, respectively (Figures 2C and 2D). Expression of H3E50K and H3E50D also significantly increases the ability of HMECDD cells to form colonies in limited dilution, clonogenic assays compared to wildtype H3.3 or H3.1 (Figures 2E-H). In fact, H3.3E50K expression yields more robust clonogenic growth than stable expression of either H3.3K27M or H3.3K36M. Moreover, the ability of these H3E50 histone variants to enhance cellular transformation from an ectopic expression vector demonstrates that H3E50K and H3E50D can function in a dominant manner, as all endogenous histone H3 genes remain wildtype and endogenous wildtype H3 is expressed (Figures 2A, 2B, Figures S2A, S2B). This result is consistent with patient data mined, in which patient tumors identified with a missense mutation that encodes H3E50K contain a single mutation in one allele of a single H3 gene (Figure 1E). Collectively, these data suggest that H3E50K, and to a lesser extent, H3E50D, exhibits growth and transformation properties consistent with a *bona-fide* oncohistone protein in both H3.1 and H3.3.

### H3E50K enhances cell migration and invasion

To determine whether H3E50 variants regulates cell migration and invasion, which are cellular phenotypes associated with cancer metastasis (50), we performed a series of cell migration assays. We cultured the series of transduced HMECDD cells to confluence, then scratched the 2D culture, and monitored the ability of cells to migrate and close the wound over time. Expression of H3.3E50K significantly increases the migration rate (Figure S3A, B) and percentage of wound closure at both 5h and 12h (Figure S3C) compared to wildtype H3.3. To further examine the propensity of H3.3E50K expression to drive cell migration, we performed transwell migration assays. H3E50K expression increases by approximately 4-fold the number of HMECDD cells that migrate through the 8.0 µm porous membrane in response to growth factors as compared to H3.3 wildtype (Figure 3A, B, Figure S3D). We also found that H3.3E50K and H3.1E50K expression enhances the invasive properties of HMECDD cells as seen by a 6-fold and 2-fold enhanced invasion through a matrigel-coated transwell membrane, respectively (Figures 3C-F, Figures S3E, F). Notably expression of H3.1E50D did not significantly increase transwell invasion compared to wildtype H3.1 (Figures 3E, F). Because the H3E50K missense mutation co-occurs with other genomic alterations, including *BRAF* mutation in human cancer (Figure 1G), we investigated whether H3E50K expression enhances oncogenic activity in the presence of co-occurring oncogenic driver events. We overexpressed wildtype H3.3 or H3.3E50K in the *BRAF* V600E-mutant melanoma cell line A2058 (Figure S3G,H). Despite the fact that H3.3E50K expression does not increase cell proliferation or clonogenic growth in *BRAF*-mutant melanoma cells (Figure S3I-K), expression of H3.3E50K with *BRAF* V600E significantly enhances cell migration and invasion in this context (Figure 3G-J, Figure S3L, M). Notably, while we detect a statistically significant difference in the ability of H3.3E50K expression to enhance proliferation and clonogenicity as well as support migration and invasion, the magnitude of the increase observed for migration and invasion is much larger, suggesting this potential oncohistone could be involved in an EMT-like process. Together, these data suggest that H3E50K expression contributes to an increase in cancer cell migration and invasion alone or in combination with known oncogenic driver gene expression in cancer types that harbor an H3E50K missense mutation.

**Figure 3.**
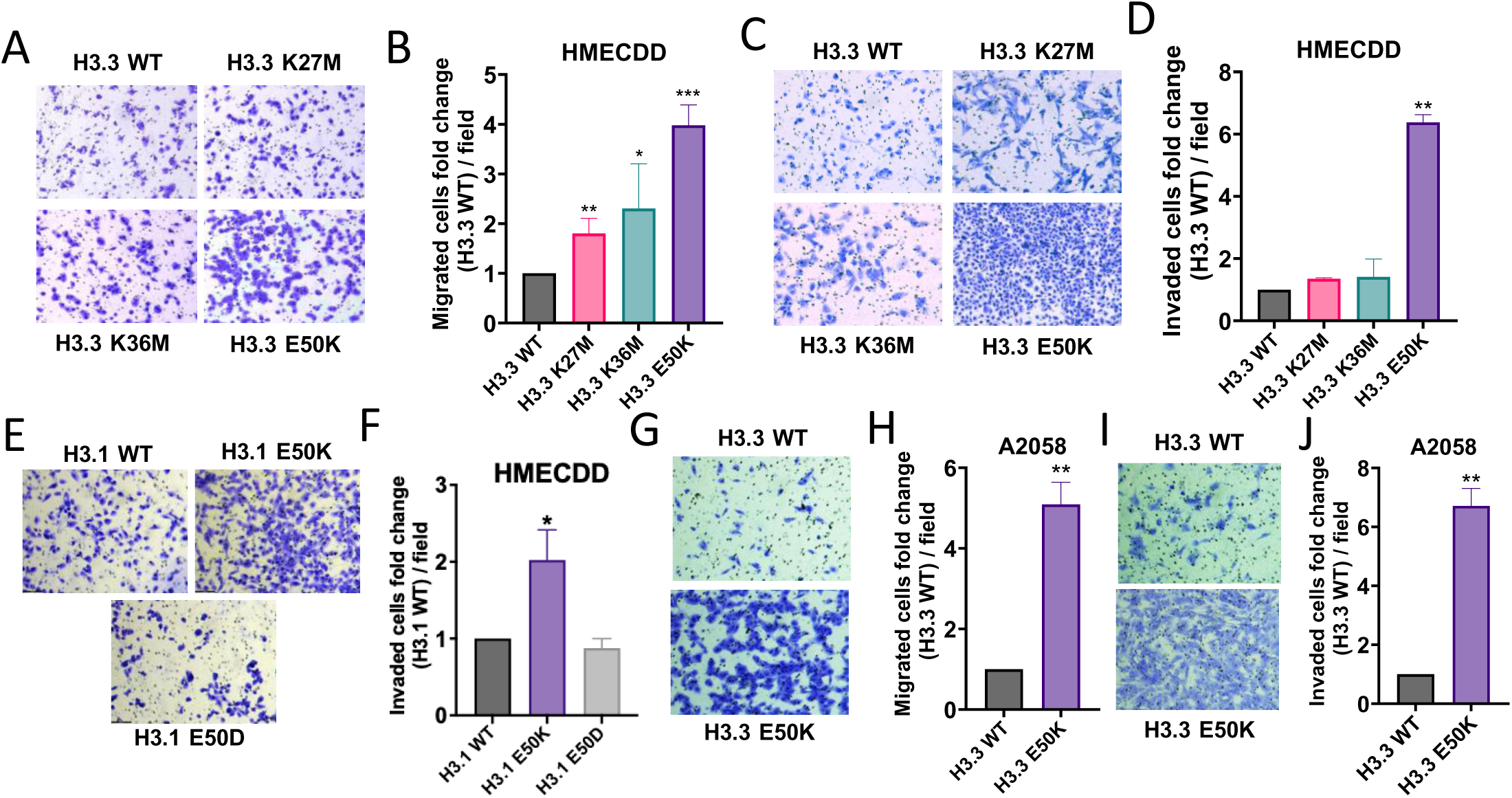
H3E50K expression enhances phenotypes associated with cancer progression. (A) HMECDD cells stably transduced with the indicated H3.3 plasmids were seeded and cell migration through an 8.0 μm filter was measured after 8 hr, representative images shown. (B) Quantification of (A), *n = 3.* (C) HMECDD cells stably transduced with the indicated H3.3 plasmids were seeded and cell invasion through an 8.0 μm 1 mg/ml matrigel-coated filter was measured after 48 hr, representative images shown. (D) Quantification of (C), *n = 2.* (E) HMECDD cells stably transduced with the indicated H3.1 plasmids were seeded and cell invasion through an 8.0 μm 1 mg/ml matrigel-coated filter was measured after 48 hr, representative images shown. (F) Quantification of (E), *n = 3.* (G) A2058 cells stably transduced with the indicated H3.3 plasmids were seeded and cell migration through an 8.0 μm filter was measured after 8 hr, representative images shown. (H) Quantification of (G), *n = 3.* (I) A2058 cells stably transduced with the indicated H3.3 plasmids were seeded and cell invasion through an 8.0 μm 1 mg/ml matrigel-coated filter was measured after 48 hr, representative images shown. (J) Quantification of (I), *n = 3*.

### H3E50K perturbs histone posttranslational modifications

Prior studies examining the molecular mechanisms by which oncohistone expression drives cancer development and progression reveal that while all known histone H3 oncohistones modulate transcriptional competence, they do so via unique mechanisms of action. To begin to explore how H3.3E50K expression could contribute to oncogenic growth, we acid extracted histones from HMECDD cells expressing wildtype H3.3-TY1, H3.3E50K-TY1, H3.3K27M-TY1, or H3.3K36M-TY1 and examined a suite of histone PTMs critical for proper regulation of gene expression. While expression of H3.3K27M dysregulates H3K27me3 and H3.3K36M dysregulates H3K36me3 on a global level, respectively, (Figure 4A, B), H3.3E50K reduces multiple PTMs, including those with opposing functions. H3.3E50K globally reduces H3K4me1 and H3K36me3, both of which are PTMs associated with transcriptional activation (51). In addition, H3.3E50K globally reduces H3K27me3, which is a mark of heterochromatin (51) (Figure 4A, B).

**Figure 4.**
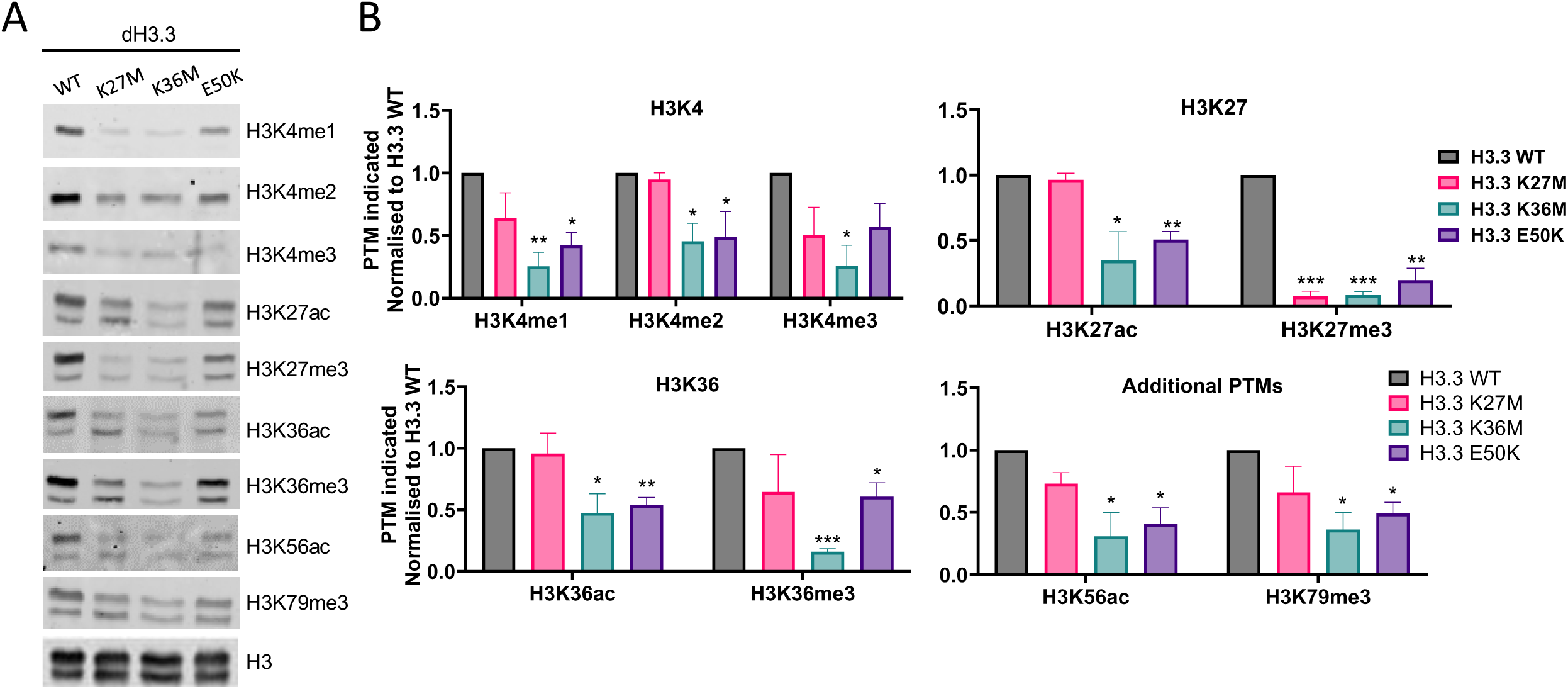
H3E50K expression perturbs chromatin modification and gene expression dynamics. (A) HMECDD cells stably transduced with the indicated plasmids and histone lysates were acid extracted as described in Materials and Methods. For each sample, 40 µg of lysates was immunoblotted with the indicated antibodies. Representative images are shown. (B) Quantification of (A) for the following modifications: H3K4, H3K27, H3K36, H3K56ac and H3K27me3, *n = 3.* All results shown are normalized to the WT H3.3, which was set to 1.0.

### H3E50K dysregulates the breast transcriptome to support EMT

Transcriptomic analysis via RNA-sequencing demonstrates that stable overexpression of H3.3E50K in HMECDD cells significantly dysregulates gene expression as compared to HMECDD cells that overexpress control wildtype H3.3 (269 increased transcripts, 85 decreased transcripts; p < 0.05) (Figure 5A, B, and Figure S4). Fast gene set enrichment analysis (FGSEA) with cancer hallmarks suggests that H3.3E50K expression positively regulates EMT, TNFα via NFκB signaling, KRAS signaling, JAK/STAT3 signaling, and other hallmarks of cancer, while decreasing hallmarks, including the G2M checkpoint (Figure 5B). Analysis of differentially expressed transcripts (Figure 5C) in H3E50K cells identify an increase in transcripts that encode proteins with defined roles in EMT, including *PREX1, FLT1* (VEGFR1), and *NLRP2* (Figure 5D). To assess whether these changes reflect an increase in EMT, we analyzed both N-cadherin and E-cadherin protein levels. We detect an increase in N-cadherin and a decrease in E-cadherin consistent with enhanced EMT (Figure 5E, F). Collectively, these data suggest that H3E50K expression supports a transcriptional program that contributes to enhanced EMT, which is consistent with our observations that H3.1 and H3.3E50K significantly increases cell migration and invasion (Figure 3).

**Figure 5.**
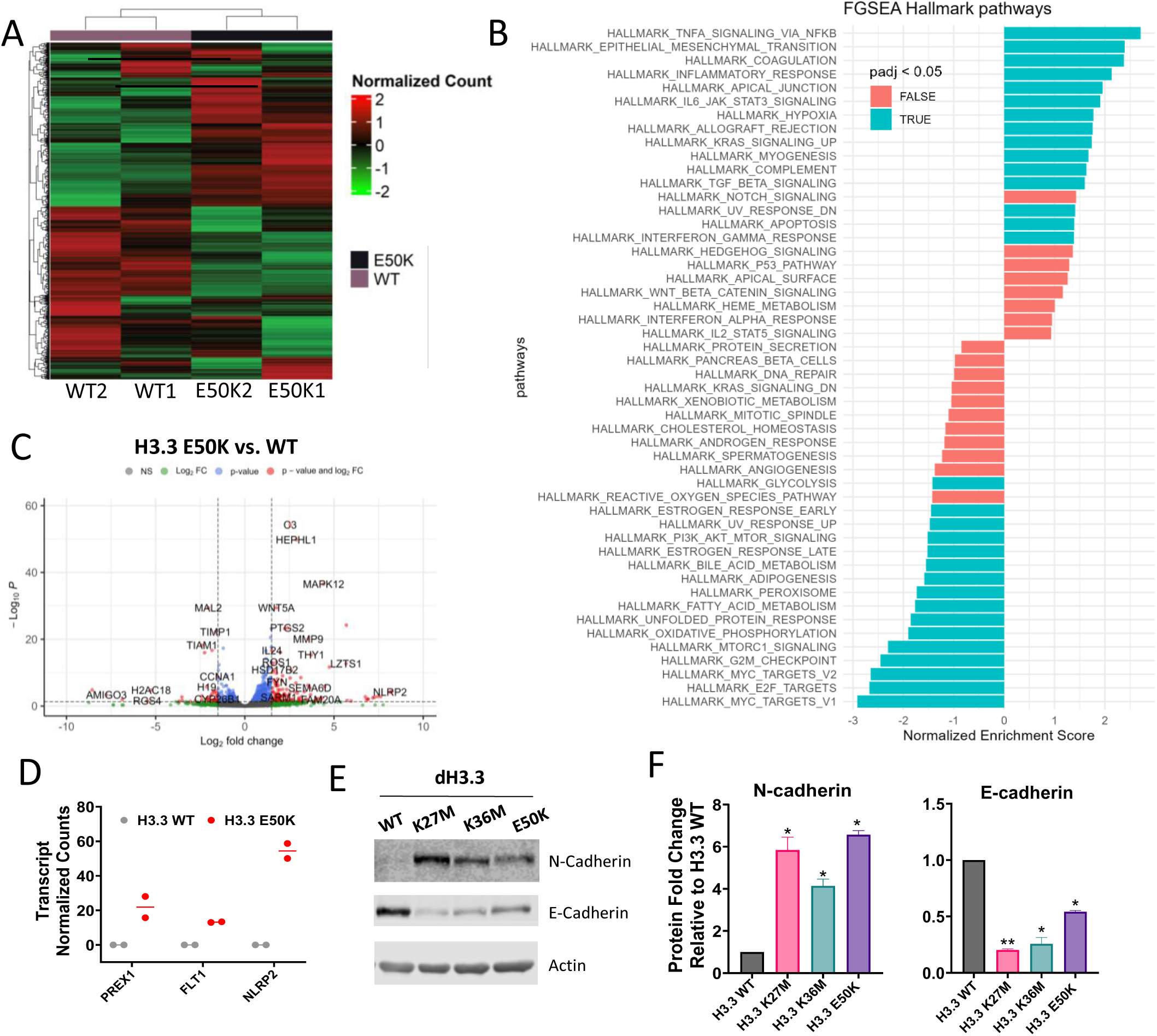
H3E50K supports an EMT gene expression program. (A) Gene cluster heatmap of HMECDD H3.3-TY1 and HMECDD H3.3E50K-TY1 cells from RNA-seq data showing differentially expressed genes. Z scores with normalized read counts were used for heat map representation. (B) FGSEA depicting upregulated or downregulated hallmark pathways from HMECDD H3.3E50K-TY1 cells compared to HMECDD H3.3-TY1 cells (blue padj < 0.05, red padj > 0.05). (C) Enhanced volcano plot depicting significantly differentially expressed genes between HMECDD H3.3-TY1 cells and HMECDD H3.3E50K-TY1 cells, where P ≤ 0.05 (blue) and log2fold change ≥ 1.5 (red). (D) Normalized read counts of indicated EMT-associated genes that are upregulated in HMECDD H3.3E50K-TY1 cells compared to H3.3-TY1. (E) HMECDD cells stably transduced with the indicated plasmids and proteins extracted. 30 µg of lysates were immunoblotted with the indicated antibodies. Representative images shown, *n = 2*. (F) Quantification of the protein levels shown in (E) normalized to actin as a loading control.

### H3E50 amino acid substitutions sensitize *S. cerevisiae* to drugs that induce various forms of cell stress

Previously characterized oncohistones cause growth defects and/or altered drug sensitivity in yeast models (7,32,34,52) demonstrating that yeast models are valuable tools for exploring cellular effects of oncohistone proteins. To explore whether H3E50 changes modeled in yeast also impact cell physiology, we utilized the model organism *S. cerevisiae.* The histone H3 proteins in budding yeast share the highest degree (89%) of identity with human H3.3 (35). As such, the nucleosome structure in *S. cerevisiae* shows similar contacts between H3E50 and H4R39, as observed in *H. sapiens* H3 (compare Figure 1G to Figure 6A). To explore the potential consequences of specific amino acid changes at this position, H3E50K, H3E50D as well as H3E50R and H3E50A were modelled both in budding yeast (Figure 6A) and human (Figure 6B) nucleosomes. As observed for H3E50K within the human nucleosome structure (Figure 1I), H3E50R creates a repulsive interaction with the coordinating H4R39 amino acid, while H3E50A eliminates the potential for interaction with H4R39.

**Figure 6.**
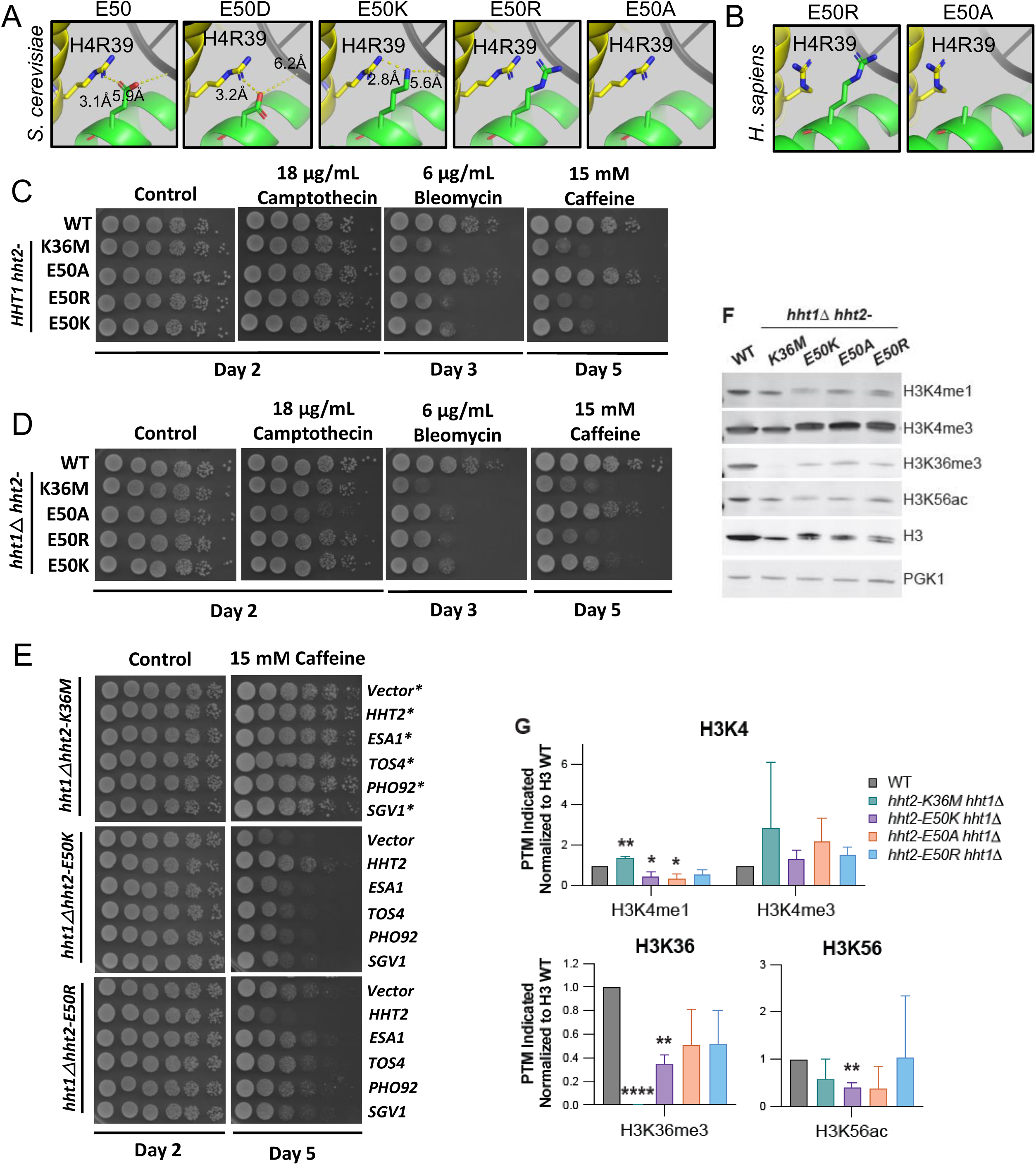
H3E50K *S. cerevisiae* exhibit restricted growth phenotypes via distinct mechanisms from H3K36M. (A) *In-silico* modeling of *S. cerevisiae* H3E50, H3E50K, H3E50D, H3E50R, and H3E50A harboring possible post-translational modifications, using PDB 1ID3 (37). (B) In-silico modeling of *H. sapiens* E50R and E50A, using PDB 5X7X (36). (C) Serial dilution spotting assays of *S. cerevisiae* cells expressing the yeast H3 homologue *HHT2* containing the indicated mutation were grown on normal media (YEPD) or media containing cellular stressors -18 µg/ml camptothecin, 6 µg/ml bleomycin, or 15 mM caffeine - for 2 to 5 days. Representative images shown, *n = 3.* (D) Serial dilution spotting assays of *S. cerevisiae* cells expressing the yeast H3 homologue *hht2* containing the indicated mutation in the absence of *HHT1* were grown on normal media (YEPD) or media containing cellular stressors -18 µg/ml camptothecin, 6 µg/ml bleomycin, or 15 mM caffeine - for 2 to 5 days. Representative images shown, *n = 3.* (E) Serial dilution spotting assays of *S. cerevisiae* cells expressing wildtype *HHT2*, *hht2-E50K, or hht2-E50R* and ectopic overexpression of known and indicated H3K36M oncohistone suppressors were grown on nutrient rich media (YEPD) or YEPD with caffeine for 2 days. Representative images shown, *n = 4.* (F) Lysates acid extracted from *S. cerevisiae* cells of the indicated genotypes were immunoblotted with the indicated antibodies. Representative images shown, *n =3.* (G) Quantification of (F). Signals were normalized to H3 and PGK1 levels.

In budding yeast, which possess two H3 genes, *HHT1* and *HHT2* (53), we engineered the *HHT2* allele to express *hht2-E50A*, *-E50R*, or *-E50K*, or, as a control, *hht2-K36M* to genetically model oncohistone expression in a background similar to oncohistone expression in human cells in which another wildtype copy of histone H3 (*HHT1*) is present. H3E50R was included to examine whether the positive charge shared between the lysine and arginine R groups impacts H3 function. Cells expressing these H3 variants were grown in control conditions or in the presence of cellular stressors or DNA damaging agents (Figure 6C). While no change in growth was detected under control conditions, *hht2-K36M*, *hht2-E50R*, and *hht2-E50K*, cells show growth defects in the presence of the DNA damaging agent bleomycin (54) as well as caffeine, a negative regulator of the TOR signaling pathway (55) (Figure 6C). These studies further demonstrate the dominant nature of H3E50K and show that H3E50R also has a dominant effect on yeast cell sensitivity to these drugs. In contrast, H3E50A does not confer a dominant growth defect.

To directly explore the functional consequences of amino acid substitutions at E50 for histone H3, we also examined yeast cell growth using mutants that express H3E50 variants as the sole copy of budding yeast histone H3. These cells express each engineered mutant *hht2* allele (*hht2-E50A*, *-E50R*, or *-E50K*) and are deleted for *HHT1*. With this experimental paradigm, the H3E50R and H3E50K cells still show slow growth in the presence of the DNA damaging agent bleomycin as well as caffeine (Figure 6D). As the sole copy of H3, H3E50A confers a growth defect in the presence of camptothecin as well as bleomycin (56), but no significant effect is detected in the presence of caffeine. Thus, converting E50 to a positive residue affects yeast cell sensitivity to specific stressors in a manner that does not occur with a neutral change, suggesting the positive charge provides a gain of function. In both genetic contexts (Figure 6C, D) H3E50R confers a slightly stronger growth defect in the presence of bleomycin or caffeine as compared to H3E50K.

### H3E50-mutant histones alter cell physiology via a mechanism that is distinct from H3K36M-mutant histones

One advantage of the budding yeast system is that simple genetics can be employed to identify cellular pathways impacted by histone PTM and/or oncohistones (34,57,58). We recently employed such an approach to identify high copy suppressors of the caffeine-sensitive growth defect of H3K36 mutants in budding yeast (34). This approach identified several chromatin modifying enzymes and complexes that when overexpressed can suppress the impaired growth of H3K36 mutants on media containing caffeine (34). As *S. cerevisiae* expressing H3E50K and H3E50R are also sensitive to caffeine (Figure 6C, D), we examined whether the growth of the H3E50 mutants like H3K36R/M mutants is suppressed by overexpression of the suppressor genes, which would suggest that similar cellular pathways are impacted in these oncohistone models. We overexpressed H3K36 suppressors, including *ESA1*, encoding a lysine acetyltransferase (59), *TOS4,* encoding a protein implicated in transcriptional regulation (60), *PHO92*, encoding an *N6*-methyladenosine (m6A) reader, (61) and *SGV1,* encoding a CDK9 kinase, in *S. cerevisiae* with H3E50K or H3E50R as the sole copy of H3 and assessed growth in the presence of caffeine (Figure 6E). Overexpression of the established H3K36 suppressors shows minimal rescue of the H3E50 mutants. Taken together, these genetic suppressor experiments suggest that H3E50 variants alter yeast cell physiology in a manner that may be distinct from H3K36M.

To determine whether H3E50K perturbs proximal PTMs in budding yeast, we acid extracted histones from the strains expressing wildtype H3 protein or mutant H3 (e.g., *hht2-E50A*, *-E50R*, or *-E50K*) that are also deleted for *HHT1.* While expression of H3K36M globally reduces H3K36me3 (Figure 6F, G), we find mutation of E50 significantly reduces H3K4me1 (Figure 6F, G), which is consistent with our findings in human models (Figure 4). Collectively, these data suggest that *S. cerevisiae* can be employed to streamline functional studies examining the molecular mechanisms of action of oncohistones, readily complementing companion studies in human systems.

## Discussion

Here, we leverage the publicly available cBioPortal and COSMIC datasets to identify histone H3E50 mutations as novel and recurrent histone mutations in human cancers. While multiple cancer-associated genomic alterations occur that result in a change of H3E50 to another amino acid, a missense mutation that changes H3E50 to a lysine to produce H3E50K is amongst the most common in these publicly available datasets. We show that ectopic expression of either H3.1H3E50K or H3.3E50K functions in a dominant manner to drive cell proliferation, growth in limited dilution, migration, and invasion. H3.3E50K also globally reduces proximal H3 PTMs involved in both transcriptional repression and activation. Transcriptomics suggest that H3.3E50K expression governs essential pathways and processes involved in cell invasion and the epithelial to mesenchymal transition, consistent with H3E50K-mediated cellular phenotypes. Genetic suppressors assays in *S. cerevisiae* demonstrate that while restricted growth in caffeine is shared between the known oncohistone H3K36M and H3E50K, genes that suppress growth phenotypes of H3K36M cells do not show similar suppression of H3E50 mutant models. Taken together, these data suggest that H3E50K may alter cell physiology and confer oncogenic properties in a manner that is distinct from previously analyzed oncohistones.

Unlike the H3K36M and H3K27M oncohistones, missense mutations that change H3E50 to other residues, including H3E50K, occur at a lower frequency without a preference for a specific tissue of origin, with H3E50K identified in breast, lung, and skin cancers, in addition to other cancer types (Figure 1F, Figure S1B). We modelled H3.1E50K and H3.3E50K expression in untransformed human mammary epithelial cells (HMECs) that were previously transduced with dominant negative p53 (HMECDD). Expression of H3E50K supports a transformed phenotype in this context. While we observed a modest but significant increase in the proliferation and clonogenic expansion of cells, the ability of H3.3E50K expression to increase cell migration and invasion is not only significant, but of a large magnitude, suggesting that H3.3E50K expression may be involved in the initiation of metastatic phenotypes. We also examined the transformative capacity of H3E50K in a second untransformed human breast cell line, MCF10A. Unlike HMECs, ectopic expression of H3E50K in the MCF10A background was insufficient to support oncogenic activity (data not shown). Because H3E50 genomic alterations occur in patient tumors in the context of other co-occurring genomic alterations (e.g., loss of function *TP53* mutation and/or activating *BRAF* V600E mutation), we investigated whether H3.3E50K expression in the *BRAF* V600E-mutant A2058 melanoma cell line enhances the transformative phenotypes already observed in A2058. While H3.3E50K expression in A2058 melanoma cells did not increase proliferation or clonogenicity, H3.3E50K expression in this context strikingly increases cell migration and invasion (Figure 3), in line with our observations in HMECDD cells. These data highlight how the overall genomic context may inform the driver nature of oncohistone mutations, including H3E50K. Without loss of p53, H3E50K expression was insufficient to drive *in vitro* oncogenic phenotypes. While H3.3E50K expression combined with the loss of function *TP53* mutation with *BRAF* V600E mutation in A2058 melanoma cells did not further increase cell proliferation, H3.3E50K did augment invasive phenotypes that are associated with EMT. While H3E50K mutation is relatively infrequent and, as such, patient outcome data are not readily available, these data raise the possibility that some oncohistone mutations, such as H3E50K, may be associated with poor patient prognosis or may be associated with later stage diagnosis due to a more aggressive phenotype, especially in situations where H3E50K co-occurs with additional oncogenic driver events. Gene signatures associated with H3E50K expression (Figure 5) support these conclusions, as H3.3E50K expression in untransformed breast cells increase transcripts associated with KRAS signaling, which could synergize with known oncogenic KRAS/MAPK signaling effectors such as *BRAF* V600E.

As described above, our findings suggest that H3.1 and H3.3E50K expression supports oncogenic activity and a transformed phenotype primarily by supporting migration and invasion in *in vitro* assays. While these experiments support the conclusion that H3E50K is a bona-fide oncohistone, *in vivo* experiments are required to assert that H3E50K mutation functions as an oncogenic driver. Moreover, our studies demonstrate H3E50K drives migratory and invasive cellular phenotypes in both untransformed HMEC cells characterized with concurrent dominant negative p53 expression, and in the context of H3.3E50K, also enhances invasion in the presence of concurrent oncogenic drivers. Transcriptional studies support these findings by demonstrating that H3E50K cells overexpress genes associated with EMT. These studies provide insight into the role H3E50K may have in cancer progression and migration as opposed to cancer initiation, suggesting that H3E50K may support cancer cell dissemination into distal sites. Conclusively testing this hypothesis in *in vivo* murine models is a critical future direction.

The previously characterized H3K27M oncohistone primarily reduces H3K27 trimethylation in *cis* and *trans* (9); similar changes are observed for H3K36M activity on H3K36 trimethylation in *cis* and *trans* (10). Additional alterations to histone PTMs may occur in these contexts, which are likely to be indirect changes as a result of the global misregulation of H3K27 methylation and H3K36 methylation, respectively. In contrast, H3.3E50K globally reduces numerous H3 PTMs associated with both transcriptional activation and repression (Figure 4).

While these data do not rule out the possibility that expression of H3.3E50K may directly modulate one or more proximal PTMs through changes in association of chromatin modifying enzymes with H3, one possible hypothesis is that H3E50K generally destabilizes the nucleosome, as has been reported for the H2BE76K oncohistone (17). Bagert, et. al biochemically examined *in situ* histone dimer exchange, and found that H3.1 E50 mutations (H3E50A/D/K/Q) enhance dimer exchange relative to wildtype H3.1(7), which suggests that H3E50K mutation may reduce nucleosome stability. Subsequent experiments should address these hypotheses and could include examination of how H3E50K perturbs the H3 interactome, and whether expression of H3E50K systematically alters the stability of nucleosomes containing H3E50K, and whether this impacts chromatin accessibility.

Our data in budding yeast demonstrate that both *hht2-E50K* and *hht2-E50R* cells are sensitive to DNA damage and TOR inhibition in the presence of wildtype endogenous histone H3, while *hht2-E50A* grows normally in the presence of all cell stressors (Figure 6C). These results align with previous studies that identified sensitivity to several DNA damaging agents in *hht2-E50A* cells (62,63). These studies both demonstrate the utility of *S. cerevisiae* as a model to functionally dissect the role of amino acid properties in contributing to oncogenic activity and suggest that the positive charge in the lysine and arginine R groups may support a pro-oncogenic phenotype (Figure 5C, D). While both *hht2-E50R* and *hht2-E50K* exhibit growth defects in the presence of the DNA damaging agent bleomycin as well as the cell stress inducer caffeine, the growth characteristics are different, suggesting that while the conserved positive charge may play a role in oncogenic activity, we cannot rule out the possibility of additional functions that differentiate H3E50K from H3E50R. These functions may enhance our understanding of why H3E50K is identified as a recurrent mutation in cancer, while H3E50R is not (Figure 1). Another study that examined *hht2-E50A* yeast cells revealed that this amino acid change can extend chronological lifespan of these cells (64), a phenotype that could contribute to oncogenic properties in human cells. In the future, to address the potential contributions of H3E50 changes, one could engineer human cell lines to express the suite of H3E50 mutant proteins and examine their propensity to support oncogenic activity and perturb gene expression, compared to H3E50K.

Current research is focused on delineating the mechanistic basis of how oncohistones, including H3K27M and H3K36M, function, with the goal to define therapeutic targets. This has led to the preclinical testing of compounds targeting a variety of transcription factors, kinases, and chromatin modifying enzymes, as well as the clinical testing of the DRD2 antagonist ONC201 in H3K27M-mutant gliomas (25,27,28). Our studies identify a series of proteins and biological processes that are upregulated, some of which may function as viable therapeutic targets for patients with cancers characterized by H3E50K. H3E50K upregulates KRAS signaling and co-occurs with *BRAF* mutation in melanoma and other cancers. As such, patient tumors characterized by H3E50K mutation may predict enhanced or prolonged sensitivity to combined BRAF and MEK inhibition via debrafinib and trametinib, respectively, which is currently a standard-of-care for the treatment of *BRAF* V600E-mutant melanomas (65). While this combination has yet to be tested in the context of H3E50K mutant cell lines and tumors, other approaches can be utilized to define H3E50K vulnerabilities. To identify functional vulnerabilities in oncohistone expressing cells, we have previously exploited *S. cerevisiae* and performed high copy suppressor screens in H3K36M/R yeast strains (34). In this context, we identified conserved chromatin modifying enzymes including the histone acetyltransferases Esa1/Tip60. This strategy can be employed in parallel with human and murine *in vitro* and *in vivo* studies to identify therapeutically actionable targets in H3E50K cell lines and tumors.

## Data Availability

The data generated in this study are publicly available in the Gene Expression Omnibus at GSE (GSE246607). Strains and plasmids are available upon request. The authors affirm that all data necessary for confirming the conclusions of the article are present within the article, figures, and tables.

## Acknowledgements

We thank the Spangle, Corbett, and Hong laboratories for helpful project discussions. We thank Dr. Anna M. Kenney for kindly providing access to the Leica Delimited microscope equipped with the MC170 camera used for imaging migration and invasion assays. We thank Dr. Dan Yan, Department of Pediatrics, Emory University for providing support for the Sartorius Incucyte.

## Author contributions

KS, CYJ, AHC, and JMS designed the research. KS, CYJ, MA, SL, RSL, JF, and SE performed experiments and analyzed the results. AHC and JMS supervised the studies. JMS wrote the manuscript with input from KS, CYJ, RSL, and AHC.

## Funding and Additional Information

This study was supported by grants from the NIH (R21CA256456 to JMS and AHC, and K12GM000680 to CYJ), the NSF (S352L5PJLMP8 to JMS), the Winship Cancer Institute of Emory University (Winship Invest$ to JMS and AHC), and Emory University Research Council (URC, to JMS and AHC).

---

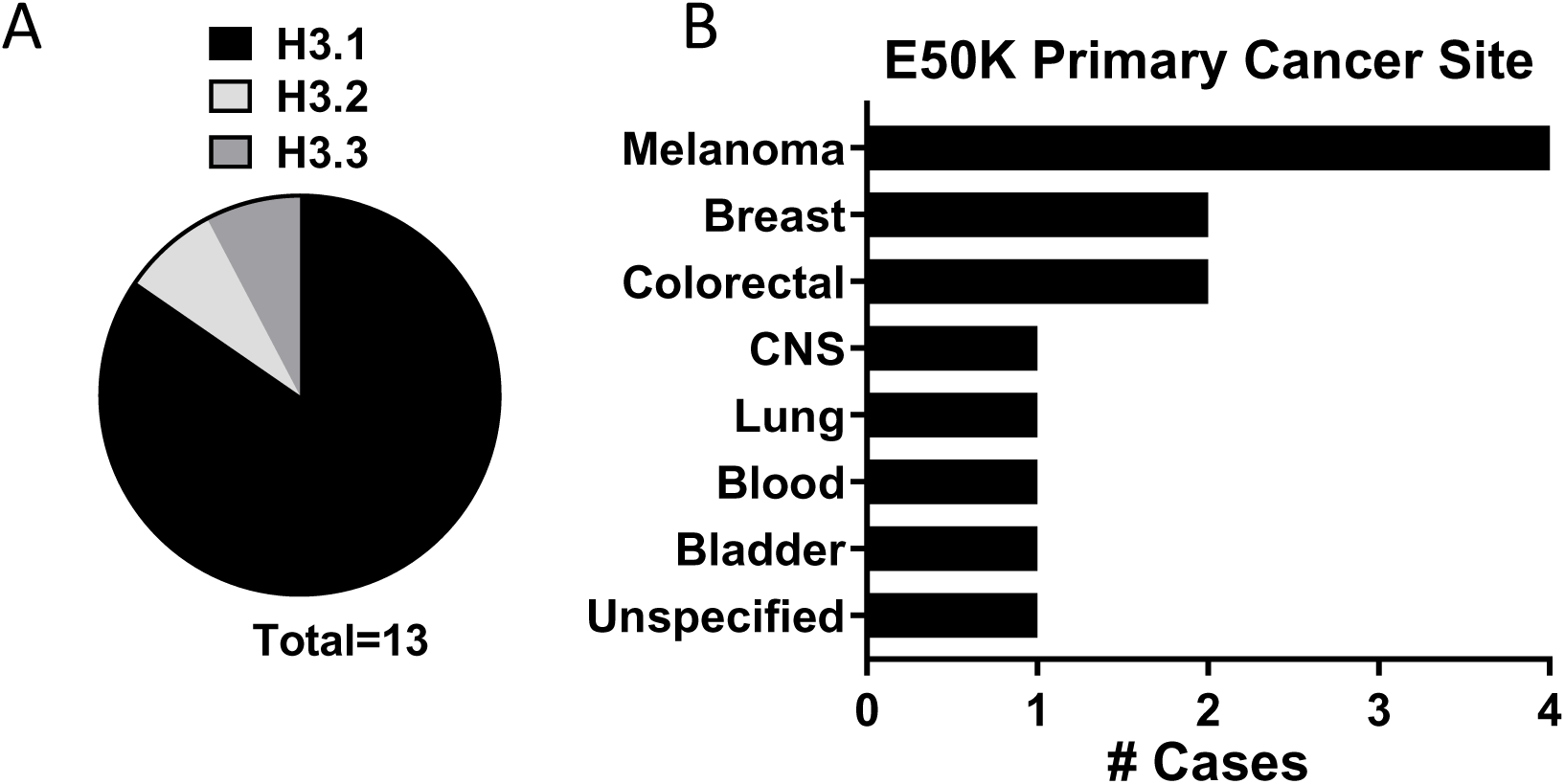

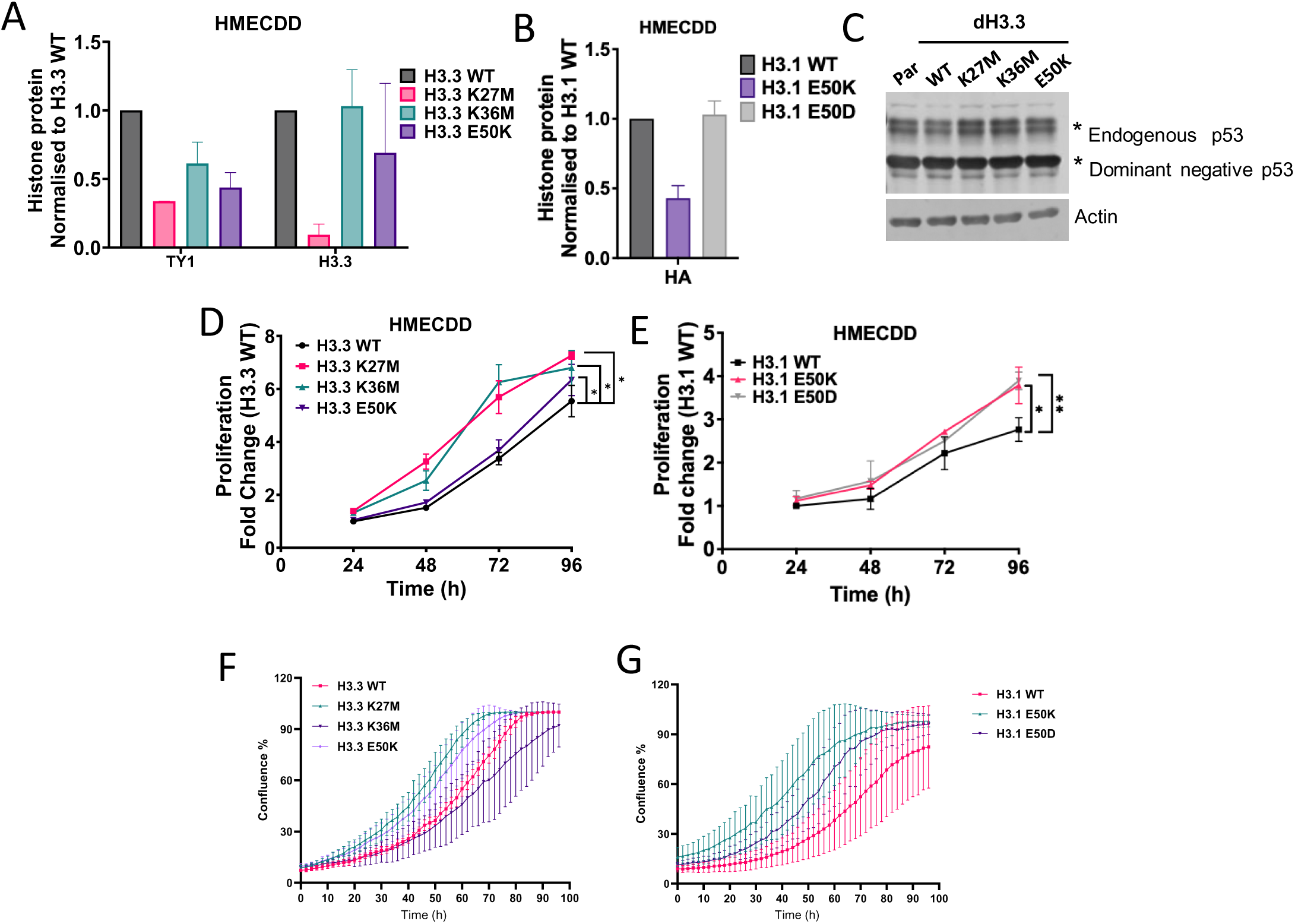

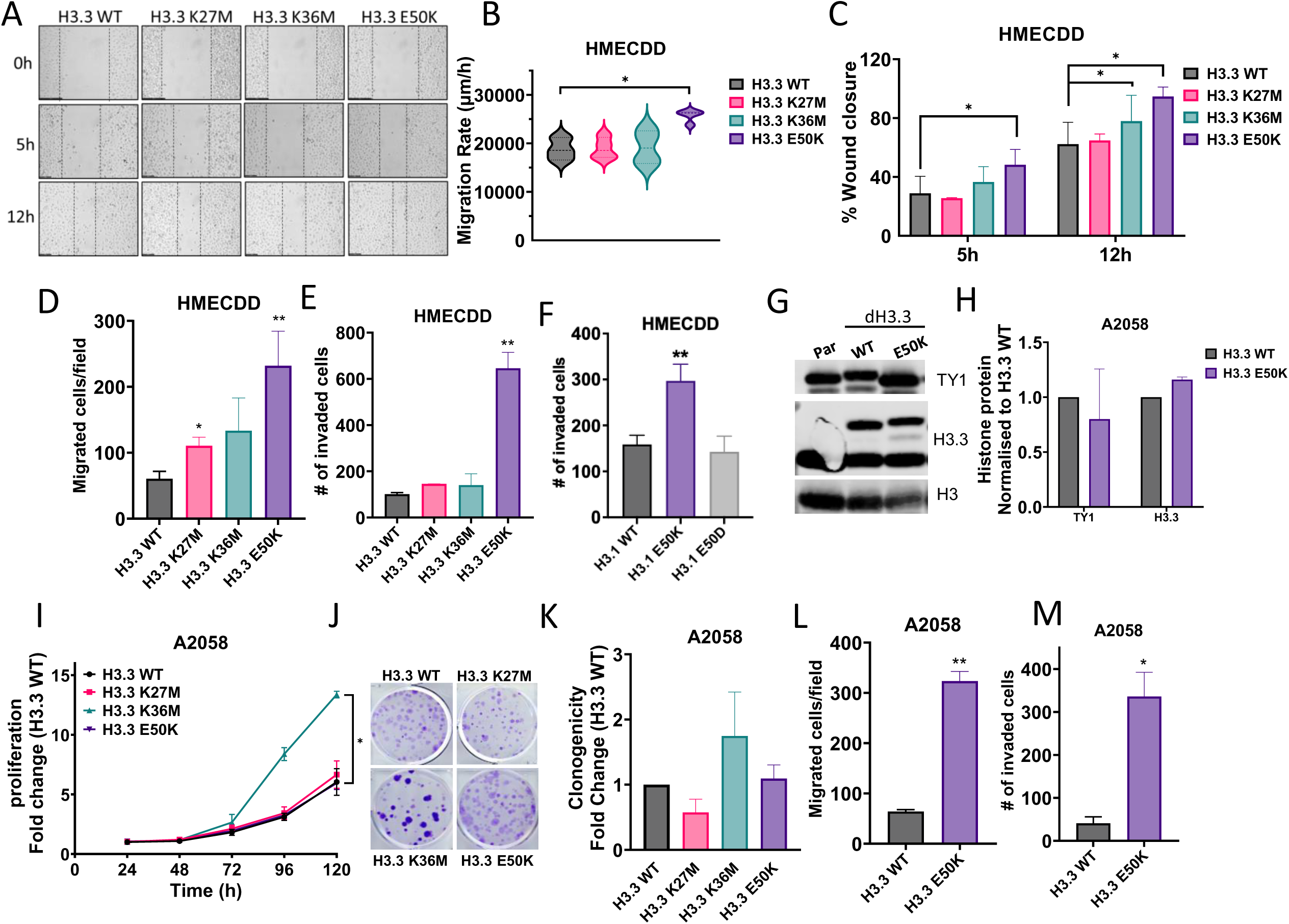

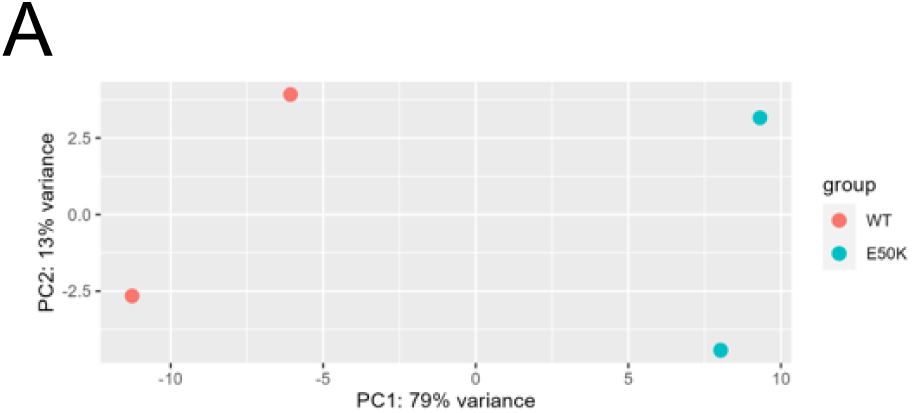

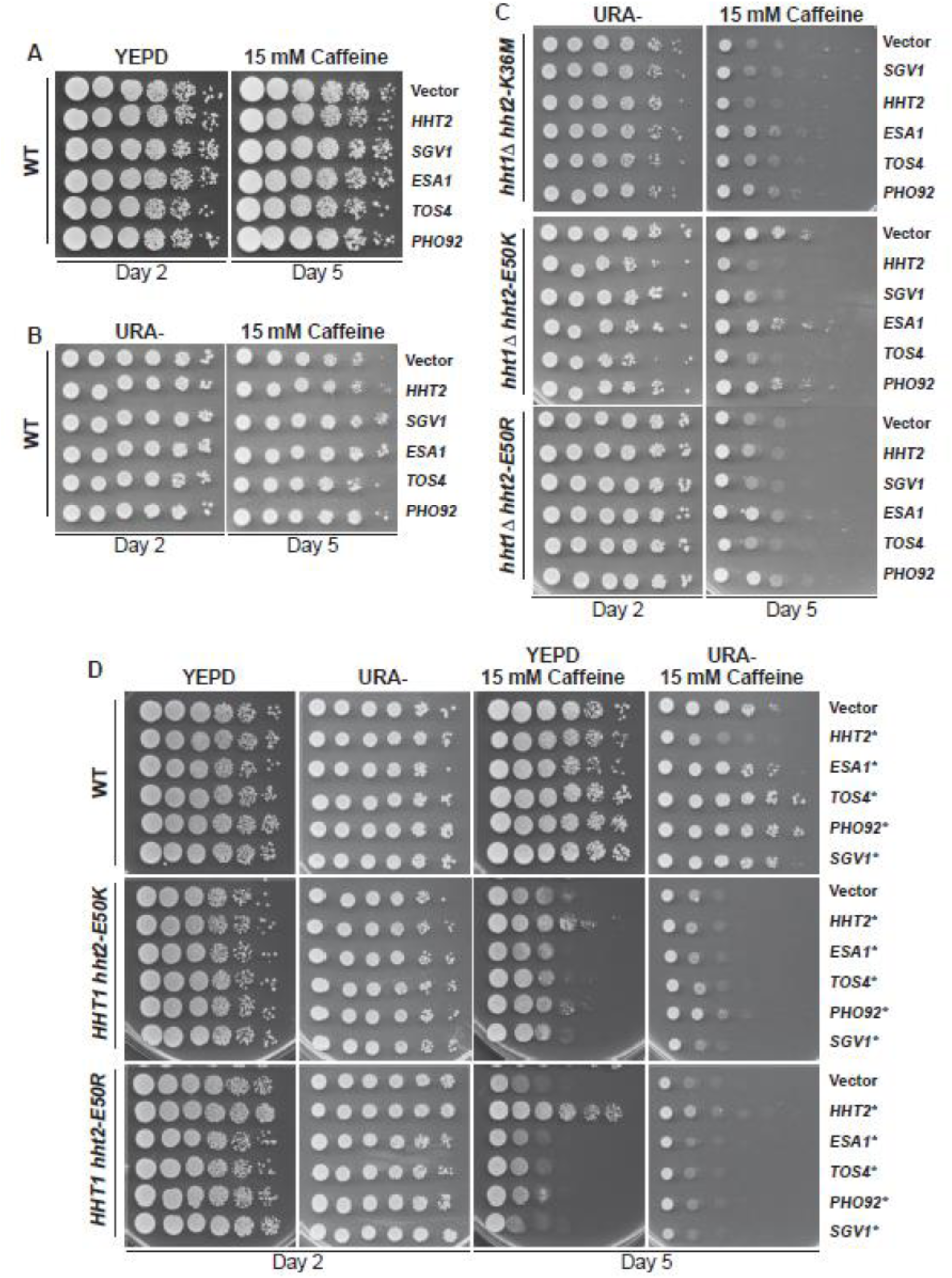

