## Supplementary Data for "Histone H3 E50K mutation confers oncogenic activity and supports an EMT phenotype"

**Supplementary Figure Legends:**

**Figure S1. H3E50K mutation occurs in human cancers.** (A) Survey of E50K mutation abundance in histone H3 genes encoded in the human genome. (B) Abundance of H3E50K mutation in human cancers.

**Figure S2. H3E50K expression enhances cancer-associated phenotypes.** (A) Quantification of H3.3 mutant expression via TY1 epitope tag and H3.3 in parental HMECDD cells, or HMECDD cells stably expressing pBabePuro H3.3-TY1, H3.3K27M-TY1, H3.3K36M-TY1, or H3.3E50K-TY1, $n = 3$. (B) A2058 cells stably transduced with pBabePuro H3.3-TY1 or H3.3E50K and lysates acid extracted. 40 µg lysates were immunoblotted with the indicated antibodies. Representative images shown, $n = 2$. (C) Quantification of H3.3 mutant expression via TY1 epitope tag and H3.3 in parental A2058 cells, or A2058 cells stably expressing pBabeP H3.3-TY1 or H3.3E50K-TY1, $n = 3$. (D) Stable A2058 cells expressing the indicated H3.3 mutant proteins were seeded, and cell proliferation measured over the indicated time course. $n = 3$. (E) A2058 cells stably transduced with the indicated H3.3 plasmids were seeded and cell clonogenicity measured after 12d, representative images shown. (F) Quantification of (E), $n = 3$. (G) representative images from wound healing assay over time. (H) Quantification of the migration rate (µm/hour) of HMECDD stable cells expressing the indicated H3.3 proteins in wound closure assays, $n = 3$. (I) Quantification of the percent of wound closure at 5h or 12h for HMECDD cells stably transduced with the indicated H3.3 plasmids, $n = 3$. (J) Bar graph for the number of migrated HMECDD cells stably transduced with the indicated H3.3 plasmids from 8 µm transmembrane filter after 8h ($n = 3$).

**Figure S3. H3E50K expression perturbs chromatin modification and gene expression dynamics.** (A) Principle component analysis (PCA) of transcriptomic data from HMECDD cells stably expressing H3.3-TY1 or H3.3E50K-TY1.

**Figure S4. Additional controls confirm that H3K36M suppressors do not rescue growth defects in H3E50 mutant cells.** (A) Serial dilution spotting assay of *S. cerevisiae* cells expressing the yeast H3 homologue *hht2* containing the indicated mutation were grown on minimal media (URA-) or media containing 15 mM caffeine plates for 2 or 5 days to select for plasmid expression. (B) Serial dilution spotting assay of wildtype *S. cerevisiae* that overexpress known and indicated H3K36M suppressors were grown on control media (YEPD) or 15mM caffeine plates for 2 or 5 days to assess whether these genes confer enhanced growth in a nonspecific manner. (C) The same wildtype and *hht2* mutant *S. cerevisiae* strains as in Fig 5E were grown on URA- control and 15 mM caffeine plates for 2 or 5 days to ensure plasmid expression. (D) The same suppressor plasmids as in Fig. 5E were transformed into wildtype or *hht2* mutant strains. These H3E50K/R strains express wildtype *HHT1*, as opposed to the *hht1*Δ strains in Fig. 5E.

Figure S1

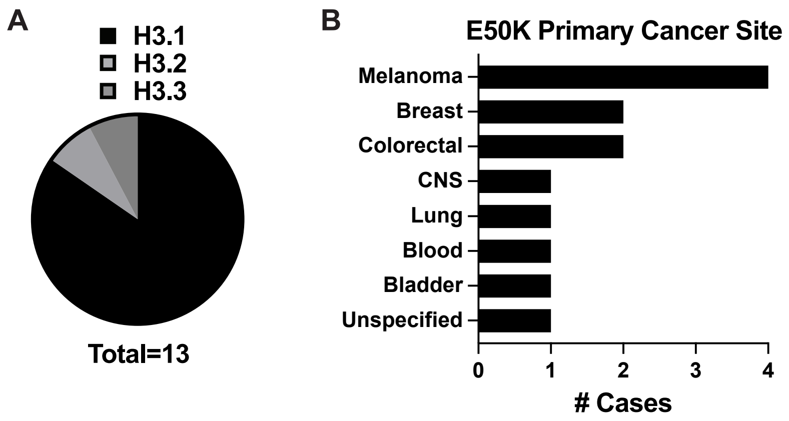
A
H3.1
H3.2
H3.3
Total=13
B
E50K Primary Cancer Site
Melanoma
Breast
Colorectal
CNS
Lung
Blood
Bladder
Unspecified
0
1
2
3
4
### Cases

Figure S2

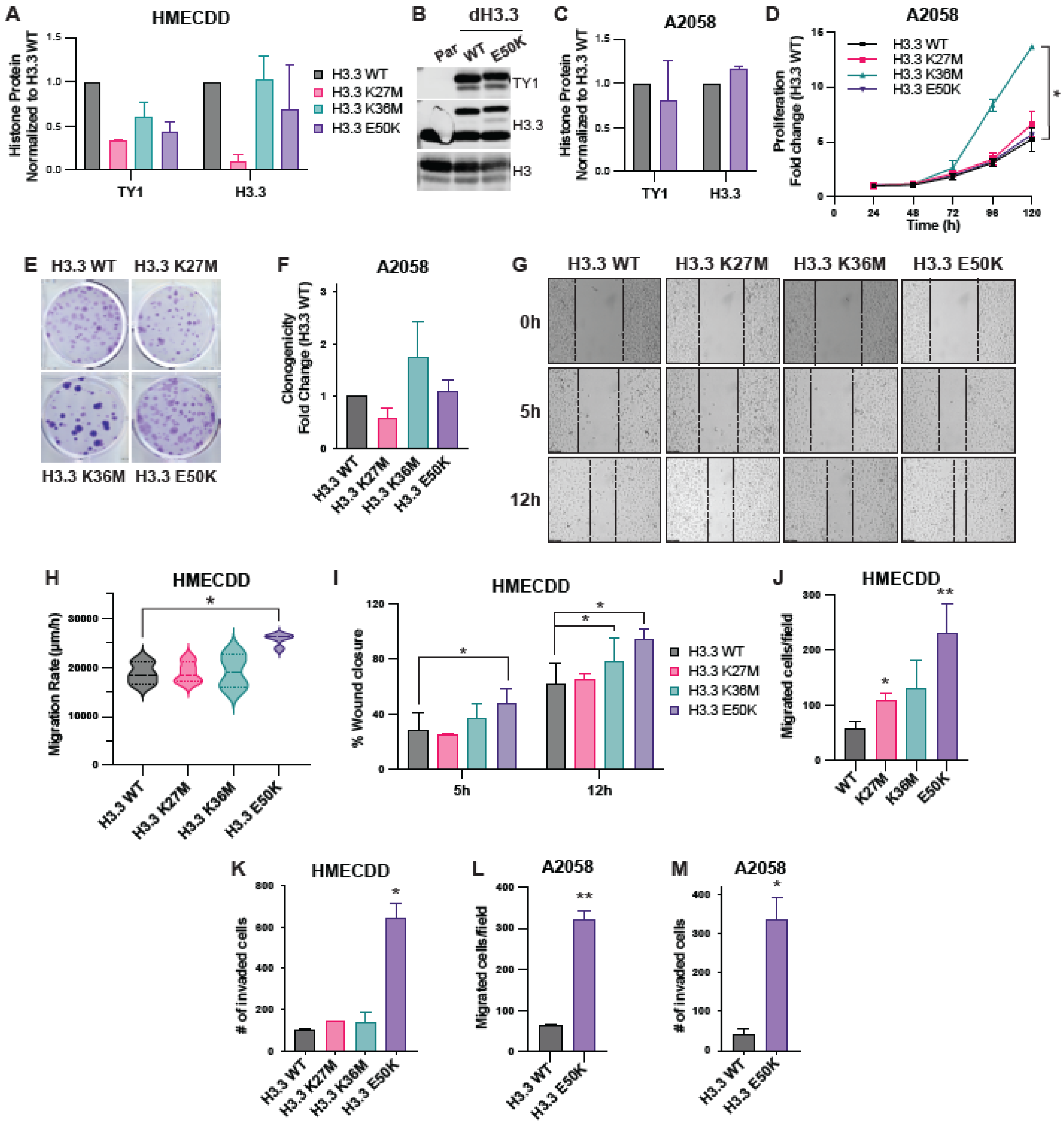

A
HMECDD
Histone Protein Normalized to H3.3 WT
TY1
H3.3
H3.3 WT
H3.3 K27M
H3.3 K36M
H3.3 E50K
B
dH3.3
Par
WT
E50K
TY1
H3.3
H3
C
A2058
Histone Protein Normalized to H3.3 WT
TY1
H3.3
D
A2058
Proliferation Fold change (H3.3 WT)
Time (h)
H3.3 WT
H3.3 K27M
H3.3 K36M
H3.3 E50K
E
H3.3 WT
H3.3 K27M
H3.3 K36M
H3.3 E50K
F
A2058
Clonogenicity Fold Change (H3.3 WT)
G
H3.3 WT
H3.3 K27M
H3.3 K36M
H3.3 E50K
0h
5h
12h
H
HMECDD
Migration Rate (μm/h)
I
HMECDD
% Wound closure
5h
12h
J
HMECDD
Migrated cells/field
WT
K27M
K36M
E50K
K
HMECDD
### of invaded cells
L
A2058
Migrated cells/field
M
A2058
### of invaded cells

Figure S3

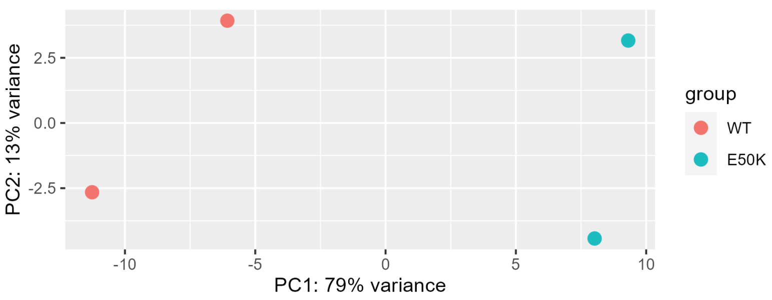

PC2: 13% variance
2.5
0.0
-2.5
-10
-5
0
5
10
PC1: 79% variance
group
WT
E50K

Figure S4

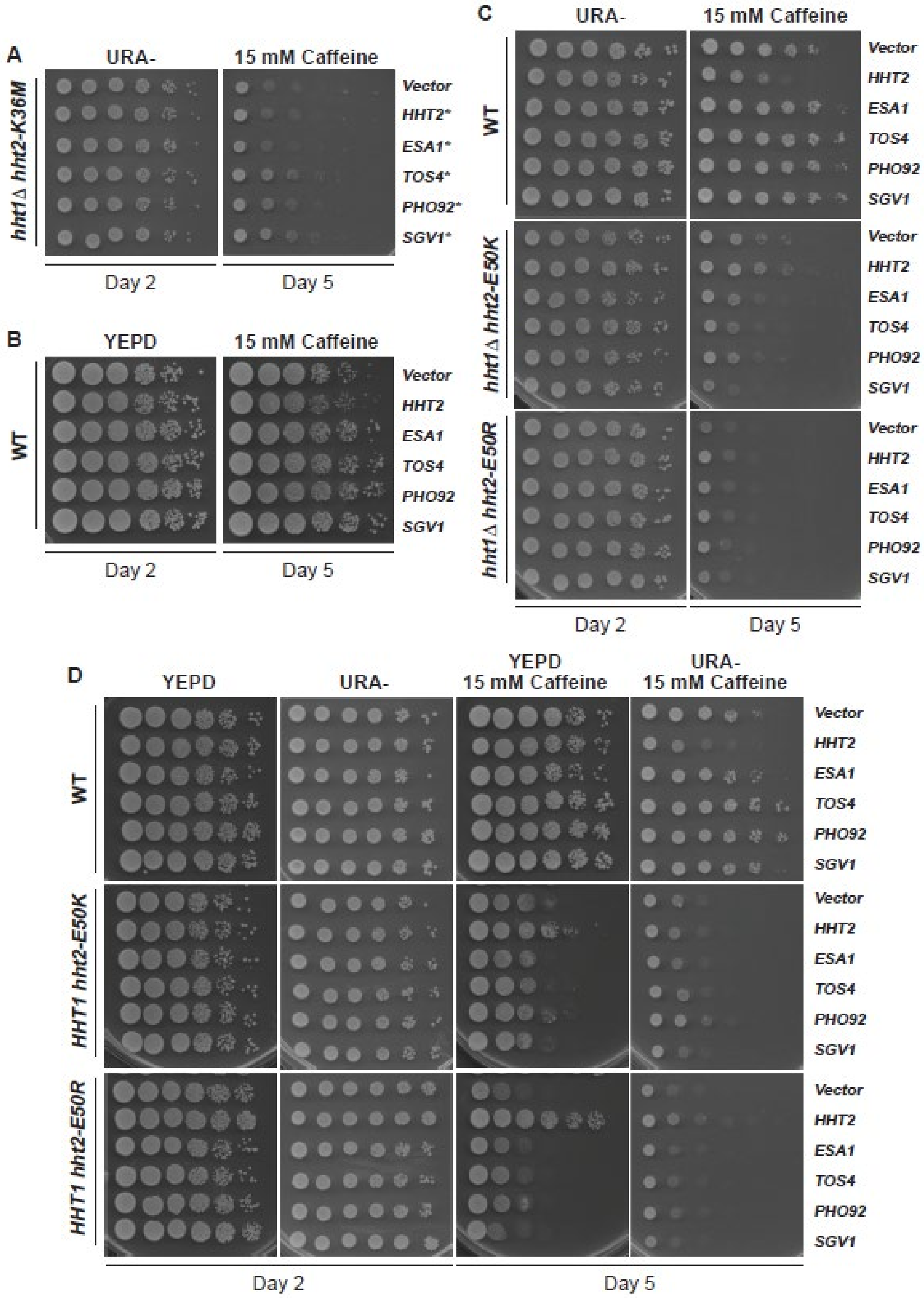

A
URA-
15 mM Caffeine
hht1Δ hht2-K36M
Vector
HHT2*
ESA1*
TOS4*
PHO92*
SGV1*
Day 2
Day 5
B
YEPD
15 mM Caffeine
WT
Vector
HHT2
ESA1
TOS4
PHO92
SGV1
Day 2
Day 5
C
URA-
15 mM Caffeine
WT
hht1Δ hht2-E50K
hht1Δ hht2-E50R
Vector
HHT2
ESA1
TOS4
PHO92
SGV1
Day 2
Day 5
D
YEPD
URA-
YEPD 15 mM Caffeine
URA- 15 mM Caffeine
WT
HHT1 hht2-E50K
HHT1 hht2-E50R
Vector
HHT2
ESA1
TOS4
PHO92
SGV1
Day 2
Day 5

**Supplementary tables and table legends:**

**Table S1. Co-occurring genomic alterations present in patient tumors harboring H3E50 mutation.** Genomic alterations for each patient were extracted from the COSMIC and cBioPortal databases and known and probable oncogene and tumor suppressor genes were filtered from the list. These data represents the co-occurring genomic alterations found in patient tumors harboring H3E50*, H3E50D, H3E50Q and H3E50K mutations.

| **Histone H3E50 alteration** | **Primary tumor site** | **Co-occurring genomic alterations in known or probable oncogenes/tumor suppressor genes** | **Amino acid change** |
|---|---|---|---|
| E50* | Colorectal | *PIK3CA* | E542K |
| | Lung | *BRCA2*<br>*KRAS*<br>*MRE11*<br>*NF1*<br>*SMAD4*<br>*TP53* | R2842C<br>Q61H<br>X642_splice<br>K476T<br>R361H<br>R213* |
| E50D | Breast | *PIK3CA*<br>*ARID1B*<br>*RB1* | N345K, E542K, H1047R<br>Q254*<br>s565*, q93* |
| | Bladder | *ARID1A*<br>*FGFR3*<br>*KDM6A*<br>*KRAS*<br>*PTEN*<br>*TP53* | Q386*<br>S249C<br>Q208*<br>E63K<br>Q245*<br>Q192* |
| | Blood | *TP53* | D189Tfs*19, |
| | HNSCC | *ATR*<br>*NOTCH*<br>*TP53* | H2559N<br>R365C<br>tp53 x187_splice, E285V_splice |
| | Liver | *ATM*<br>*CTNNB1* | R337H<br>D32N |
| | lung | *SETD2*<br>*SMAD4*<br>*TP53* | S512*, E325*<br>Y353C<br>n239D |
| E50Q | Breast | *FOXP1*<br>*PIK3CA*<br>*PTEN* | Q182*<br>E545K, H1047R<br>N48I |
| | Bladder | *CREBBP*<br>*KDM6A*<br>*TP53* | E594*<br>D1062*<br>E249S |
| | Blood | *TP53* | X125_splice |
| | Cervical | *TP53* | R248W |
| E50K | Breast | *ATR*<br>*NOTCH4*<br>*PIK3CA*<br>*SMAD4* | E650K<br>R385C<br>H1047R, E545K<br>A459T |

| | | | |
|---|---|---|---|
| | Bladder | *FGFR3* | Y373C |
| | | *PIK3CA* | V165I |
| | | *SMARCB1* | Q368* |
| | Blood | *BRAF* | G469A |
| | | *STAT3* | D661A |
| | Colorectal | *BRCA1* | E907K |
| | | *ESR1* | N519S |
| | | *KRAS* | K117N |
| | | *PIK3CA* | M1043I, H1047I |
| | | *PTEN* | E299* |
| | | *SMAD4A* | I530L |
| | | *TP53* | K132T |
| | Lung | *NF1* | Q83* |
| | | *STK11* | I303M |
| | | *TP53* | E349* |
| | Melanoma | *BRAF* | P764S, P804S, V600K |
| | | *BRCA2* | Q1063* |
| | | *NF1* | R440*, G629R |
| | | *NOTCH4* | V1712L, |
| | | *TP53* | P386I |

**Table S2. Yeast strains and plasmids used in this study.**

| Strain/Plasmid | Description | Source |
|---|---|---|
| Wildtype (yADP127) | *MATa; his3Δ200 leu2Δ1 ura3-52 trp1Δ63 lys2-128delta* | Duina and Turkal 2017 (1) |
| *hht2*-K36M *hht1Δ* (ACY 2822) | *MATa; his3Δ200 leu2Δ1 ura3-52 trp1Δ63 lys2-128delta; hht2-K36M; hht1Δ::KanMX* | Lemon *et al*. 2022 (2) |
| *hht2*-E50A *hht1Δ* (ACY 2964) | *MATa; his3Δ200 leu2Δ1 ura3-52 trp1Δ63 lys2-128delta; hht2-E50A; hht1Δ::KanMX* | This Study |
| *hht2*-E50R *hht1Δ* (ACY 2968) | *MATa; his3Δ200 leu2Δ1 ura3-52 trp1Δ63 lys2-128delta; hht2-E50R; hht1Δ::KanMX* | This Study |
| *hht2-E50K hht1Δ* (ACY 2972) | *MATa; his3Δ200 leu2Δ1 ura3-52 trp1Δ63 lys2-128delta; hht2-E50K; hht1Δ::KanMX* | This Study |
| *hht1Δ* (ACY2818) | *MATalpha; his3Δ200 leu2Δ1 ura3-52 lys2-128delta; hht1Δ::KanMX* | Lemon *et al*., 2022 (2) |
| YEp352 | *URA3*; 2μ; *amp*$^{R}$ | Hill *et al*., 1986 |
| *HHT2* (pAC4201) | *HHT2*, *URA3*, 2μ, *amp*$^{R}$ | Lemon *et al*., 2022 (2) |
| *ESA1* (pAC4190) | *ESA1*, *URA3*, 2μ, *amp*$^{R}$ | Lemon *et al*., 2022 (2) |
| *TOS4* (pAC4196) | *TOS4*, *URA3*, 2μ, *amp*$^{R}$ | Lemon *et al*., 2022 (2) |
| *PHO92* (pAC4193) | *PHO92, URA3*, 2μ, *amp*$^{R}$ | Lemon *et al*., 2022 (2) |
| *sgv1-Δ2+8aa* (pAC4208) | *sgv1-Δ2+8aa*, *URA3*, 2μ, *amp*$^{R}$ | Lemon *et al*., 2022 (2) |
| *HHT2_HHF2* from screen (pAC4145) | *HHT2*, *HHF2*, *URA3*, 2μ, *amp*$^{R}$ | Lemon *et al*., 2022 (2) |
| *SGV1* from screen (pAC4132) | *SGV1*, *URA3*, 2μ, *amp*$^{R}$ | Lemon *et al*., 2022 (2) |
| *ESA1* from screen (pAC4149) | *ESA1*, *URA3*, 2μ, *amp*$^{R}$ | Lemon *et al*., 2022 (2) |
| *TOS4_YLR184W* from screen (pAC4150) | *TOS4*, *URA3*, 2μ, *amp*$^{R}$ | Lemon *et al*., 2022 (2) |
| *PHO92_WIP1_BCS1* from screen (pAC4160) | *PHO92*, *WIP1*, *BCS1*, *URA3*, 2μ, *amp*$^{R}$ | Lemon *et al*., 2022 (2) |
